# Integrative analysis reveals extensive interactions among C2H2 zinc finger proteins at chromatin loop anchors

**DOI:** 10.64898/2026.03.06.710197

**Authors:** Ernest Radovani, Edyta Marcon, Syed Nabeel-Shah, Shuye Pu, Guoqing Zhong, Hua Tang, Hongbo Guo, Irene M. Kaplow, Andrew Emili, Timothy R. Hughes, Jack F. Greenblatt

**Affiliations:** Donnelly Centre, University of Toronto, Toronto, ON, M5S 3E1, Canada; Department of Molecular Genetics, University of Toronto, Toronto, ON, M5S 1A8, Canada; Department of Computer Science, Stanford University, Stanford, CA 94305, USA; Department of Biological Sciences, Carnegie Mellon University, Pittsburgh, PA 15213, USA; Ray and Stephanie Lane Department of Computational Biology, Carnegie Mellon University, Pittsburgh, PA 15213, USA; Department of Biochemistry, Oregon Health Sciences University, Portland, OR 97239, USA; OHSU Knight Cancer Institute, Portland, OR, USA

## Abstract

Cys2-His2 -zinc-finger proteins (C2H2-ZFPs) form the largest class of human transcription factors, yet their potential role (s) in higher-order genome organization has remained largely poorly defined. To date, only a small number of family members, e.g., CTCF and YY1, have been shown to function in chromatin organization. Here, we examined the global relationship between C2H2-ZFPs and long-range chromatin interactions (LRIs). By integrating ChIP-seq datasets for 216 C2H2-ZFPs and genome-wide maps of chromatin looping, we observe that more than 40% of the human C2H2-ZFPs are significantly enriched at LRI anchors. We found that depletion of a subset of such C2H2-ZFPs is associated with altered expression of genes linked by the LRIs they occupy. To investigate whether protein-protein interactions (PPIs) contribute to this pattern, we generated a large-scale interactome using affinity purification followed by mass spectrometry experiments, encompassing 345 C2H2-ZFPs, and LUMIER binary interaction assays for 204 C2H2-ZFPs. We identified 1,732 binary interactions, suggesting extensive connectivity among the C2H2-ZFP family members. Integrative analysis of PPI, ChIP-seq, and chromatin interaction datasets revealed that interacting C2H2-ZFP pairs are significantly co-enriched at LRIs and frequently localize to either the same or opposing loop anchors. Finally, by correlating ChIP-seq with cancer mutational datasets, we observe that DNA-binding sites of ∼35% of LRI-associated C2H2-ZFPs overlap somatic mutations in cancer genomes. Together, our results reveal a widespread network of C2H2-ZFP interactions associated with chromatin loop anchors, providing an important resource for elucidating mechanisms regulating chromatin organization.

## Introduction

Transcription factors (TFs) bind specific DNA sequences in gene regulatory elements and can recruit cofactors to regulate gene expression (1). In humans, more than 1,600 DNA-binding proteins have been identified, which can be grouped into families based on the structures of their DNA-binding domains (1, 2). Among these, C2H2-zinc-finger proteins (C2H2-ZFPs) constitute the largest class of TFs, with over 700 members (1, 2). Most C2H2-ZFPs follow a similar domain architecture, where they contain multiple zinc finger domains (ZnFs), up to 40 per protein, with a median of nine, towards their C-termini that enable recognition of long and information-rich DNA motifs (3, 4), since each ZnF can bind 3 nucleotides (5). Many C2H2-ZFPs also harbour auxiliary domains towards their N-termini that are thought to participate in protein-protein interactions. The three most notable auxiliary domains include the KRAB (Krüppel-associated box), SCAN (SRE-ZBP, CTfin51, AW-1, and number 18 cDNA), and BTB/POZ (BR-C, ttk, and bab/Pox virus and zinc finger) domains, with KRAB being the most abundant; these domains can also be used to define major subfamilies of the C2H2-ZFPs (3). The KRAB domain interacts with the corepressor TRIM28 (also known as KAP1) and is commonly associated with transcriptional repression (6, 7). The SCAN and BTB domains are thought to dimerize with other SCAN and BTB domains, thus permitting the formation of homo- and/or hetero-dimers (8–12).

We and others have previously used ChIP-seq and ChIP-Exo to identify DNA-binding sites for a large subset of the C2H2-ZFPs (13–15). These studies revealed that C2H2-ZFPs bind diverse DNA sequence motifs that are generally consistent with *in silico* predictions based on the amino acid sequences of their zinc fingers (13, 14, 16). KRAB C2H2-ZFPs often bind endogenous repeat elements and recruit TRIM28 to repress such elements (13–15, 17). Non-KRAB C2H2-ZFPs, on the other hand, are often associated with more open chromatin, and they frequently interact with functional elements such as promoters and enhancers (14).

Three-dimensional chromatin organization is a basic feature of eukaryotic genomes that is highly regulated and has been increasingly shown to play important roles in gene regulation (18, 19). A small number of C2H2-ZFPs have also been implicated in the formation of chromatin long-range interactions (LRIs) (20–23). For example, CTCF, YY1, and SP1 form LRIs by mechanisms that involve self-association or interaction with Cohesin - a protein complex that is thought to extrude chromatin and facilitate loop formation (21, 22, 24–28). More recently, MAZ, another C2H2-ZFP, has been reported to interact with CTCF during loop formation (23, 29). Such protein interactions can juxtapose distal regulatory elements (DREs) with promoters, thereby influencing transcriptional output (30). However, whether association with LRIs represents a broader property of the C2H2-ZFP family, and whether protein-protein interactions (PPIs) among C2H2-ZFPs play any role in it, remain largely unexplored.

Affinity purification coupled to mass spectrometry (AP/MS) has been widely used to map interactomes across diverse organisms and infer protein function(31–41). Previous AP/MS studies of subsets of C2H2-ZFPs revealed their interaction with a diverse set of nuclear factors involved in transcription, chromatin, and post-transcriptional regulation (14, 42). Although these studies provided important functional insights, they surveyed less than 20% of the family and therefore did not capture the global architecture of C2H2-ZFP interaction networks.

Here, we integrate genome-wide DNA binding data, chromatin interaction maps and large-scale protein-protein interaction analyses to investigate the functional interplay between C2H2-ZFPs and long-range chromatin interactions. We report that over 40% of human C2H2-ZFPs are significantly enriched at anchors of LRIs connecting promoters and putative DREs, and that depletion of a subset of these factors is associated with altered expression levels of genes linked by these interactions. We also generated AP/MS data for 225 C2H2-ZFPs and, by combining it with our previously published protein interaction datasets, constructed a PPI network encompassing approximately half of all human C2H2-ZFPs. Our analysis reveals extensive interactions among the members of this family of TFs. Using the binary interaction assay LUMIER, we further expand this network to 1,732 interactions. Integrative analysis of PPIs, ChIP-seq binding profiles, and chromatin interaction maps reveals that interacting pairs of C2H2-ZFP frequently co-localize at the same or opposing anchors of LRIs, with multiple interacting pairs converging on individual chromatin interactions. Together, these results define a widespread network of C2H2-ZFP interactions associated with chromatin loop anchors and provide a systems-level framework for understanding how this large TF family relates to higher-order genome organization and transcription regulation.

## Results

### Widespread enrichment of C2H2-ZFPs at chromatin loop anchors

Previous studies have shown that LRIs connect promoters with DREs and can play a role in transcription regulation (43). Since certain C2H2-ZFPs, e.g., CTCF, have been shown to be critical for LRIs formation, we wanted to examine whether other family members also have a role in chromatin looping. We initiated our analysis by first defining LRIs in HEK293 cells. We analyzed RNAP II ChIA-PET (Chromatin Interaction Analysis by Paired-End Tag Sequencing) datasets generated by ENCODE and identified a high-confidence set of reproducible intrachromosomal interactions (44–46). Specifically, we analyzed over 600 million paired-end reads from the two available replicates and identified >600,000 significant intrachromosomal LRIs in each replicate that were connected by at least 2 paired-end tags (PETs) (Supplemental Fig. 1A). Among these, 63,334 LRIs were reproducible across replicates, and hence were used as high-confidence LRIs for the downstream analyses (Supplemental Fig. 1B). When we focused on interactions in which one anchor overlaps a protein-coding gene promoter and the other maps to a putative DRE, we obtained a set of 35,151 promoter-DRE LRIs with a median genomic span of ∼26 kb (Fig. 1A, Supplemental Fig. 1C).

**Figure 1.**
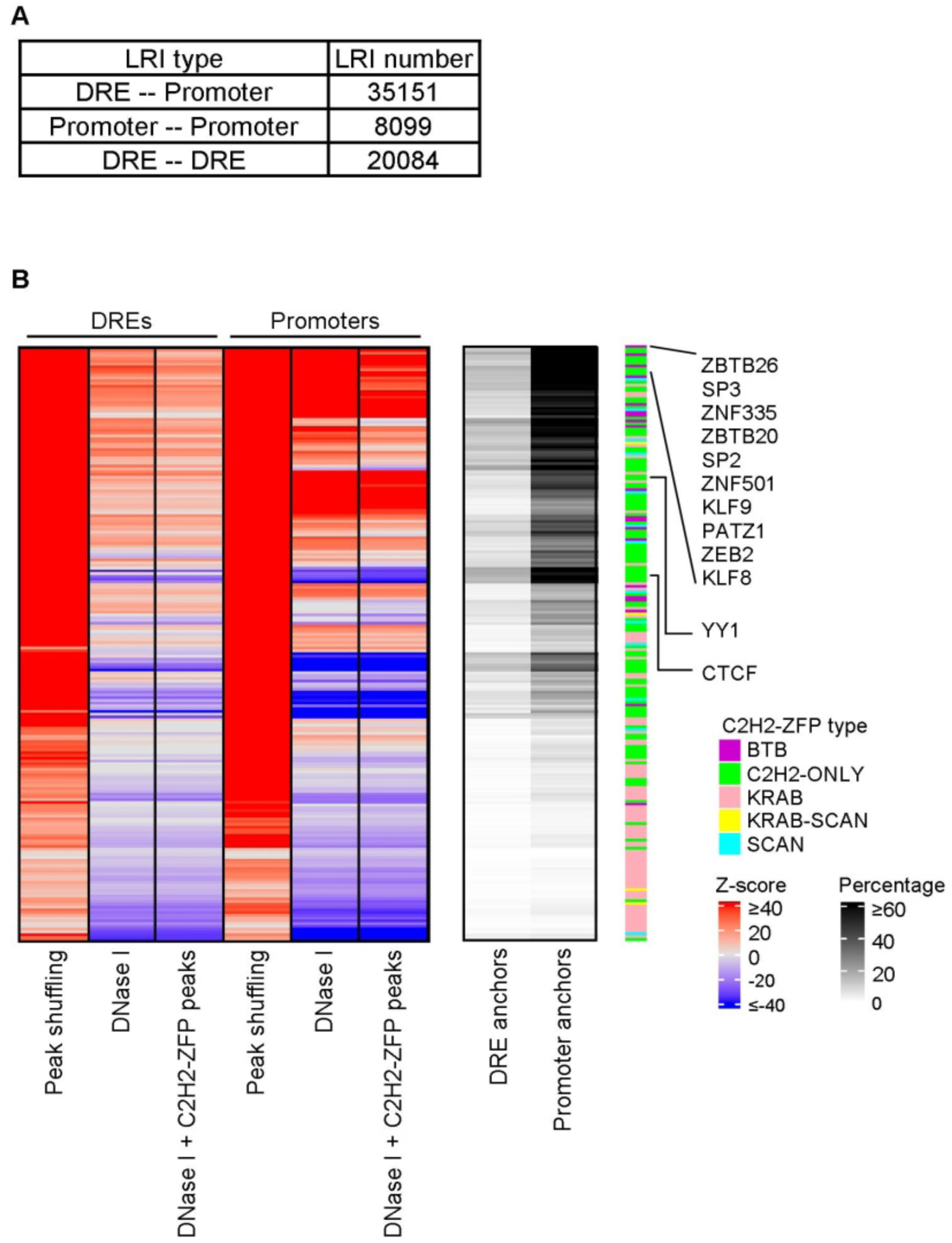
DNA-binding by C2H2-ZFPs overlaps significantly with LRI anchors. (A) Table listing the different types of LRIs that were identified from RNAP II associated ChIA-PET. (B) Heatmaps showing Z-scores for DNA-binding of C2H2-ZFPs across LRI anchors that overlap DREs or promoters (left), as well as the percentage of anchors that were bound (right). The overlaps were done using three different null backgrounds for the C2H2-ZFP ChIP-seq datasets, which consisted of 1) a random chromosome-matched set of locations for the C2H2-ZFP ChIP-seq peaks (Peak shuffling), 2) HEK293 DNase I hypersensitive sites (DNase I), and 3) HEK293 DNase I hypersensitive sites along with the ChIP-seq peaks of all 216 C2H2-ZFPs analyzed in this study (DNase I + C2H2-ZFP peaks). Only significant Z-scores are displayed.

Given the known role of certain family members in chromatin looping (20, 23, 43), we examined whether other C2H2-ZFPs are enriched at the identified LRI anchors. To investigate this possibility, we integrated publicly available ChIP-seq data for 216 C2H2-ZFPs with the chromatin interaction maps identified above (13, 14, 45, 46) (Supplemental Table S1). Specifically, we overlapped the DNA-binding sites of C2H2-ZFPs with LRI anchors that either overlap with promoters (promoter anchors) or do not overlap with promoters (DRE anchors). Using multiple background controls, we found that approximately 40% of C2H2-ZFPs are significantly enriched at either promoters or distal anchors of LRIs (Fig. 1B; Supplemental Table S2). This group includes the known chromatin loop-associated factors CTCF and YY1 (21, 43), as well as many C2H2-ZFPs not previously implicated in chromatin looping. Notably, we found that C2H2-ZFPs preferentially associate with promoter anchors, i.e., 46/216 C2H2-ZFPs bind at least 50% of promoter anchors that are involved in LRIs, whereas none of the C2H2-ZFPs bind more than 21% of the DRE anchors (Fig. 1B; Supplemental Table S2). These results suggested that association with chromatin loop anchors is a widespread property of C2H2-ZFP family members.

### Association of C2H2-ZFPs with LRIs correlates with gene expression changes upon depletion

Given that certain C2H2-ZFPs, e.g., CTCF, YY1 and MAZ, have been shown to affect gene expression via LRIs (21, 23, 47), we examined whether binding of additional C2H2-ZFPs at LRI anchors similarly affects the transcriptional output. To test this hypothesis, we analyzed publicly available RNA-seq datasets from HEK293 cells in which 16 C2H2-ZFPs had been individually depleted and examined the overlap between their DNA binding sites and LRIs linked to genes whose expression was altered upon knockdown (Supplemental Table S1). We observed that the binding sites of several of these C2H2-ZFPs at either promoter or distal LRI anchors significantly overlap with genes that were differentially expressed following depletion. Moreover, we found that individual C2H2-ZFPs appear to exhibit distinct patterns depending on whether they occupy promoter anchors, distal anchors, or both, consistent with context-dependent effects on gene expression. For example, MAZ, SP1, and ZBTB7A significantly overlap with affected genes they bind at both promoter and distal anchors, as well as at either anchor type alone (Figure 2; p-value ≤ 0.005; Fisher’s exact test). In contrast, CTCF, ZBTB48, and ZNF770 significantly (p-value ≤ 0.05; Fisher’s exact test) affect only the expression of genes that they bind at both DRE and promoter LRI anchors (Figure 2).

**Figure 2.**
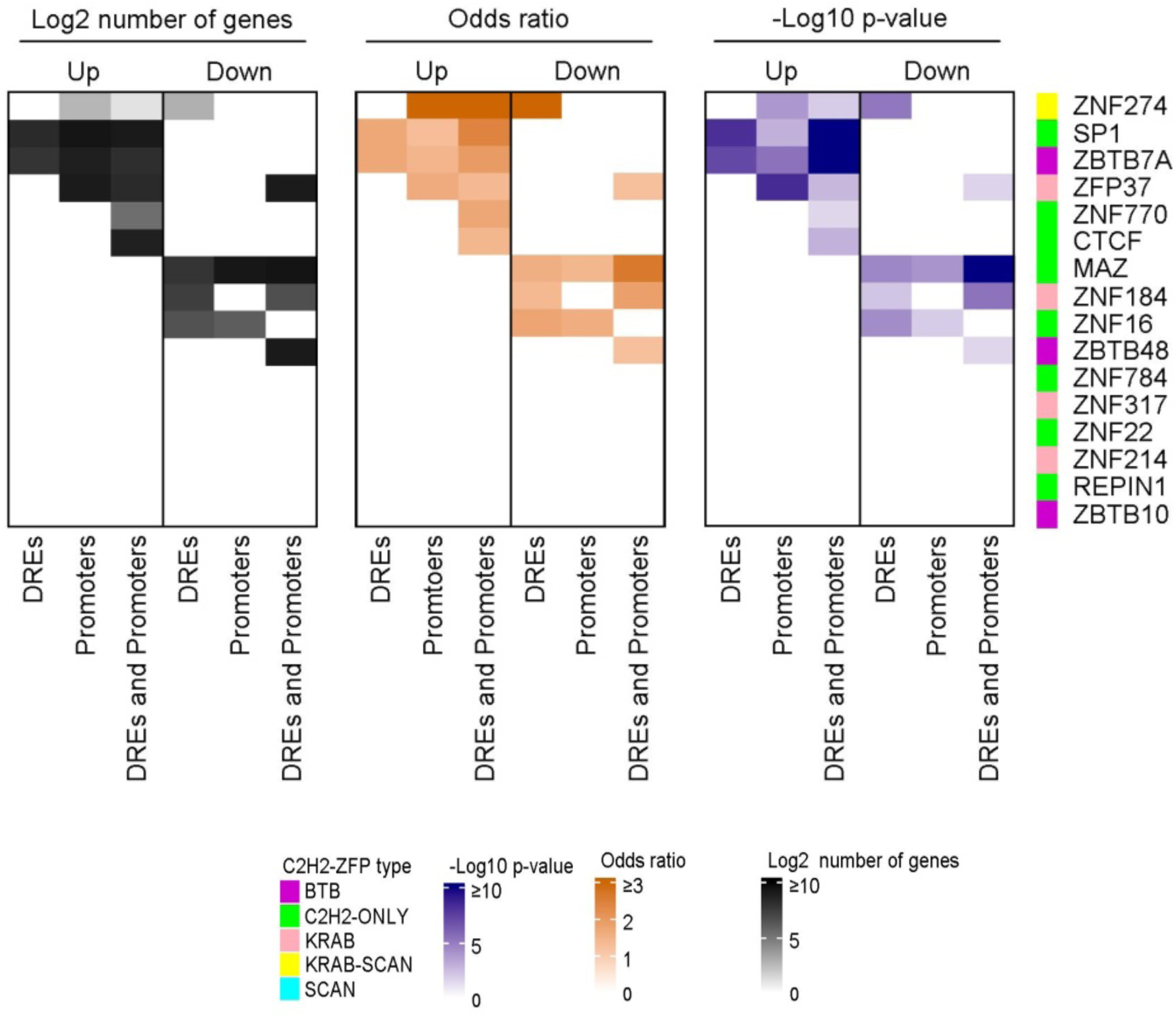
Heatmaps describing the relationship of the binding of C2H2-ZFPs at LRIs anchors to their effect on gene expression. Heatmaps indicate the Log2 number of genes that are either upregulated (Up) or downregulated (Down) following the knockdown of the C2H2-ZFPs, as well as the –Log10 p-values and the odds ratios for the association of genes that changed in expression following the knockdowns of C2H2-ZFPs with the genes that were bound by the C2H2-ZFPs at their DRE-promoter LRI anchors. The associations were analyzed separately for genes that are bound by C2H2-ZFPs at 1) only the DRE LRI anchors (DREs), 2) only at the promoter LRI anchors (Promoters), and 3) at both the DRE and promoter LRI anchors (DREs and Promoters). p-values were calculated with Fisher’s exact test, and they were corrected using the Benjamini-Hochberg method.

In almost all cases, the DNA binding of C2H2-ZFPs significantly overlap with anchors of genes that are differentially expressed following their depletion. For example, SP1 and ZBTB7A significantly overlap with anchors of genes that are upregulated (p-value ≤ 0.005; Fisher’s exact test), whereas MAZ and ZNF16 significantly overlap (p-value ≤ 0.05; Fisher’s exact test) with anchors of genes that are downregulated (Figure 2). While KRAB-containing C2H2-ZFPs are usually considered to be transcriptional repressors (7), our analysis suggests that certain KRAB-containing C2H2-ZFPs, e.g., ZNF184 and ZNF274, might be involved in the expression of genes that they bind at their DRE anchors (Figure 2). Notably, ZNF274 exhibits opposite effects depending on whether it is bound to promoter or distal anchors, i.e., promoter LRI-bound genes are upregulated, whereas downregulated genes are bound at their putative DREs. Similarly, ZFP37, another KRAB-containing C2H2-ZFP, binds to promoter anchors of genes that are upregulated following its depletion but overlaps with both upregulated and downregulated genes when binding at both anchor types. Together, these analyses suggest that C2H2-ZFPs are frequently associated with transcriptional regulation through LRIs and that their effects can be dependent on their binding configurations across interacting regulatory elements.

### A large-scale protein–protein interaction network of C2H2-ZFPs

TFs often regulate gene expression by recruiting cofactors onto chromatin through protein-protein interactions (1). To gain insight into molecular partners of C2H2-ZFPs that might be recruited at LRIs, we constructed a large-scale PPI network in HEK293 cells. After inducing the expression of GFP-tagged C2H2-ZFPs, we performed affinity purification followed by mass spectrometry (AP/MS). Of note, cell lysates were pre-treated with Benzonase, a promiscuous nuclease, to minimize DNA/RNA-mediated interactions. We generated AP/MS data for 225 C2H2-ZFPs and combined these with our previously published data for 120 additional C2H2-ZFPs (14). This yielded protein interaction data for 345 C2H2-ZFPs, representing roughly half of all human C2H2-ZFPs (Supplemental Fig. S2A; Supplemental Tables S1). These data represent the largest C2H2-ZFP PPI network to date and include representative C2H2-ZFPs from almost all C2H2-ZFP subfamilies (Supplemental Fig. S2A; Supplemental Table S3).

To provide statistical rigor, we filtered our proteomics data using ‘Significance Analysis of Interactome’ (*SAINTexpress*), an algorithm for scoring AP/MS data (48), and employed 28 GFP-alone purifications, 78 non-C2H2-ZFP TFs from various families, and 9 chromatin-related proteins as background controls (see Methods; Supplemental Fig. S2B; Supplemental Table S1). We intentionally included non-C2H2-ZFP TFs as background controls to increase the stringency of our AP/MS filtering strategy and minimize the detection of nuclear background proteins. Application of *SAINTexpress* identified 4,382 PPIs involving 333 C2H2-ZFPs and 1,808 unique prey proteins (Fig. 3A; Supplemental Table S4).

**Figure 3.**
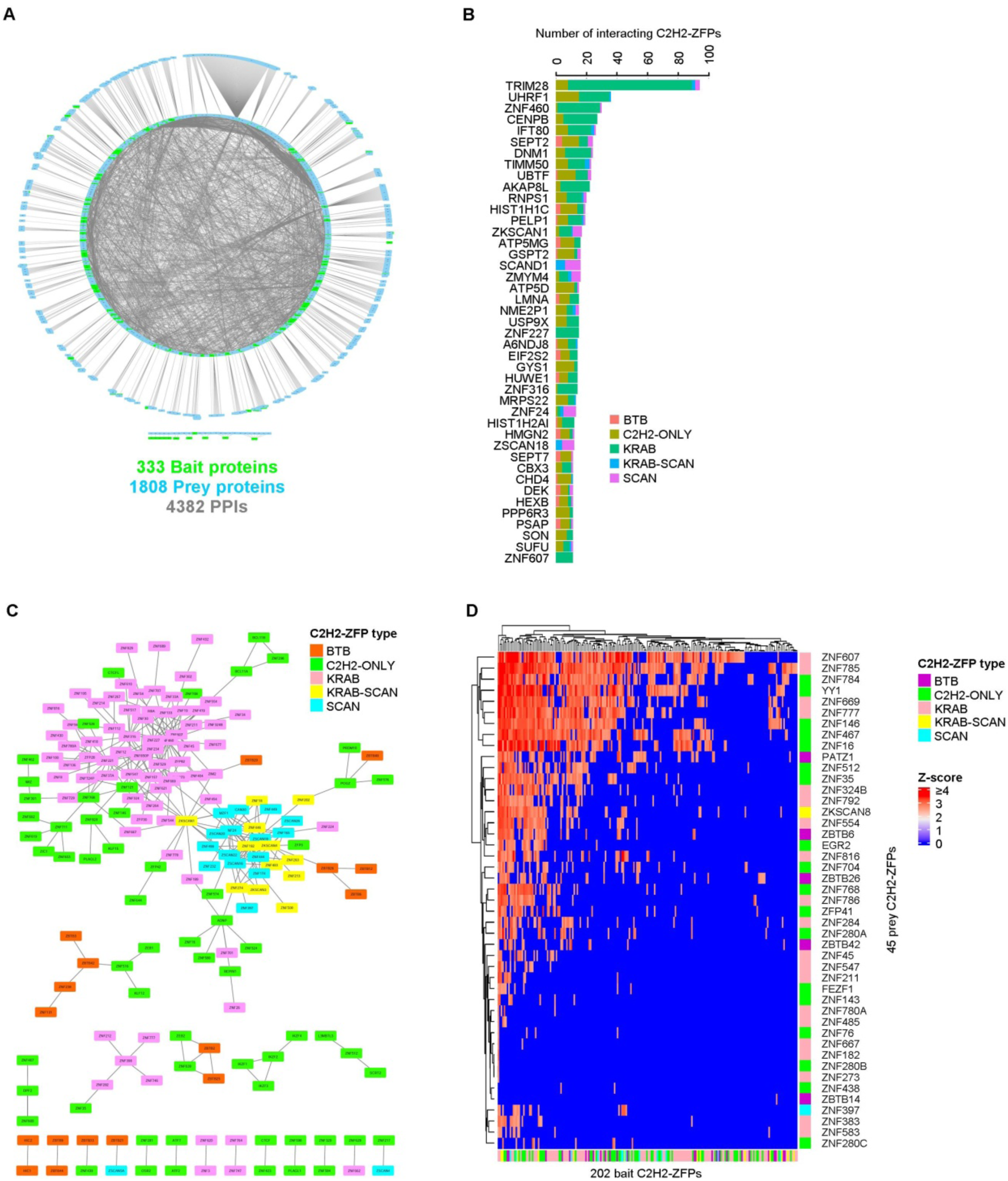
Large-scale AP/MS analysis of C2H2-ZFPs. (A) PPI network of C2H2-ZFPs (SAINT score ≥ 0.93). Bait C2H2-ZFPs are represented by green nodes, whereas interacting prey proteins are represented by blue nodes. An interactive version of this network can be found in Supplemental Data S1. (B) Barplot showing the top 43 prey proteins most frequently interacting with C2H2-ZFPs. (C) PPI network for interactions among C2H2-ZFPs obtained from AP/MS data following filtering of potential artifacts. An interactive version of this network can be found in Supplemental Data S1. (D) Heatmap showing the interactions among C2H2-ZFPs that were identified by LUMIER.

As expected, we observed that most KRAB-containing C2H2-ZFPs interacted with TRIM28, and numerous interactions were observed within the SCAN and BTB subfamilies (Fig. 3B; Supplemental Fig. S2C). While a large fraction of proteins appeared uniquely in the network, suggesting that C2H2-ZFPs mostly have distinct protein interaction profiles, certain interactors were also frequently detected. These included known transcriptional cofactors such as TRIM28, UHRF1 and PELP1, as well as proteins with less established roles in transcription, e.g., IFT80, DNM1 and ATP5L (Fig. 3B) (7, 49–52). These data suggest that C2H2-ZFPs form functional interactions with diverse proteins in cells and might participate in a range of gene regulatory pathways.

We then functionally annotated the identified prey proteins using DAVID (53). As expected, this analysis revealed strong enrichment for transcription-related proteins, consistent with the roles of C2H2-ZFPs as TFs, in addition to clusters associated with chromatin regulation, RNA metabolism and DNA repair (Supplemental Tables S5 and S6). Moreover, C2H2-ZFP interaction partners also included proteins associated with mitochondrial and metabolic processes, raising the possibility that C2H2-ZFPs may participate in diverse cellular pathways or perhaps recruit multifunctional cofactors to chromatin. We then extended our analysis to C2H2-ZFP subfamilies, finding distinct interaction profiles (Supplemental Tables S6 and S7). For example, our subfamily-specific analyses revealed that PPIs related to BTB-C2H2-ZFPs are enriched for proteasome- and neurodegeneration-associated proteins (Supplemental Fig. S2D), whereas KRAB and C2H2-only prey proteins are enriched for RNA-related factors and the KRAB-SCAN and SCAN C2H2-ZFPs PPIs are significantly enriched for Mitochondria and Transcription-related terms, respectively (Supplemental Fig. S2E-H). To gain insight into whether the evolutionary history of the C2H2-ZFPs is associated with the observed patterns, we performed a similar functional enrichment analysis for prey proteins of C2H2-ZFPs from different age groups (Supplemental Tables S5 and S6). Strikingly, prey proteins associated with the 96 and 130 evolutionarily younger C2H2-ZFPs, which are primarily restricted to great apes and primates, respectively, are depleted for transcription-related functional categories and are instead significantly enriched for RNA-associated proteins (Supplemental Fig. S2I, J). Prey proteins of C2H2-ZFPs in primates are also significantly enriched for proteins associated with Mitochondrial terms (p-value ≤ 0.05; Hypergeometric test) (Supplemental Fig. S2J). In contrast, these patterns are not observed among prey proteins of the 104 older C2H2-ZFPs (Supplemental Fig. S2K). These results indicate that C2H2-ZFPs engage diverse protein networks that may vary across subfamilies and evolutionary age groups.

### Extensive homotypic and heterotypic interactions among C2H2-ZFPs

Analysis of our AP/MS network remarkably revealed 392 interactions among 280 C2H2-ZFPs (Supplemental Table S8), suggesting that C2H2-ZFP family members extensively interact with each other. After removing interactions potentially arising from peptide misassignment or experimental carryover during MS (see Methods), 279 high-confidence C2H2-ZFP - C2H2-ZFP interactions remained (Fig. 3C; Supplemental Fig. S3A, B; Supplemental Table S8). These interactions include extensive connectivity within the KRAB-containing C2H2-ZFP and C2H2-only subfamilies, in addition to the expected interactions among SCAN- and BTB-containing proteins (8–12).

To test for interactions among C2H2-ZFP family members in an independent way, we performed a binary interaction screen using LUMIER, which can test for individual pairwise interactions of co-expressed proteins in cells (54, 55). We examined 13,850 pairwise combinations among 282 bait and 54 prey C2H2-ZFPs from various subfamilies in HEK293 cells. After filtering the data for false positives, 1,732 high-confidence interactions among 200 bait and 44 prey proteins were detected (Fig. 3D; Supplemental Fig. S3C; Supplemental Table S9). The median number of interactions per protein was eight, with a maximum of 148 for ZNF785 (Supplemental Fig. S3D), suggesting that certain C2H2-ZFPs might act as interaction hubs. Of the 74 AP/MS interactions that were tested, 21 were confirmed by LUMIER, significantly more than expected by chance (∼9 interactions) (p-value ≤ 0.0005; hypergeometric test) (Supplemental Fig. S3E). Conversely, the AP/MS data could also be used as a validation for LUMIER, although ∼20% of the LUMIER interaction partners are expressed at low levels in HEK293 cells (i.e., TPM < 5) (Supplemental Fig. S3F), and the detection of low abundance proteins can be penalized by mass spectrometry (56). This reciprocal validation using expression-filtered AP/MS data confirmed more LUMIER interactions than expected by random overlap (p-value (Supplemental Fig. S3G). ≤ 0.0005; hypergeometric test)

Together, these analyses suggested that interactions among C2H2-ZFP family members are widespread and that this interconnectivity extends beyond proteins containing SCAN and BTB dimerization domains. As such, interactions were also observed among C2H2-ZFPs lacking auxiliary domains, suggesting that zinc finger arrays and intrinsically disordered regions (IDRs) may also participate in C2H2-ZFP associations, consistent with emerging models of TF interaction networks that implicate IDRs (56).

### Interacting C2H2-ZFPs co-localize at chromatin loop anchors

Given the prevalence of interactions among C2H2-ZFPs, we examined whether the interacting C2H2-ZFPs also co-localize at chromatin loop anchors. We integrated PPI data with ChIP-seq profiles for C2H2-ZFPs and assessed co-binding at LRI anchors in two complementary ways. First, we quantified the degree of overlap *in cis* between ChIP-seq peaks of interacting and non-interacting C2H2-ZFP pairs at individual anchors. Specifically, we compared ChIP-seq co-localization at LRI anchors for 607 interacting pairs identified by LUMIER, encompassing 97 proteins with available ChIP-seq data, with all non-interacting pairwise combinations among the same proteins (Supplemental Table S10). Co-binding *in cis* (referred to as ‘degree of overlap *in cis*’ [DOVC]) was quantified as the fraction of overlapping ChIP-seq peaks at LRI anchors relative to the total peaks at LRI anchors for each pair (see Methods; Supplemental Fig. S4A; Supplemental Table S10). Interacting pairs exhibit significantly higher DOVC values than non-interacting controls (Fig. 4A), a result that remained significant when compared with an equivalent number of randomly sampled negative pairs (i.e., non-interacting C2H2-ZFPs) (Supplemental Fig. S4B). Similar trends are observed for interacting pairs identified by AP/MS using ChIP-exo datasets (Supplemental Fig. S4C). Of note, since ChIP-seq data were available for only 37 interacting pairs of C2H2-ZFPs detected by AP/MS-, we employed ChIP-exo datasets (15) to increase the statistical power in these analyses.

**Figure 4.**
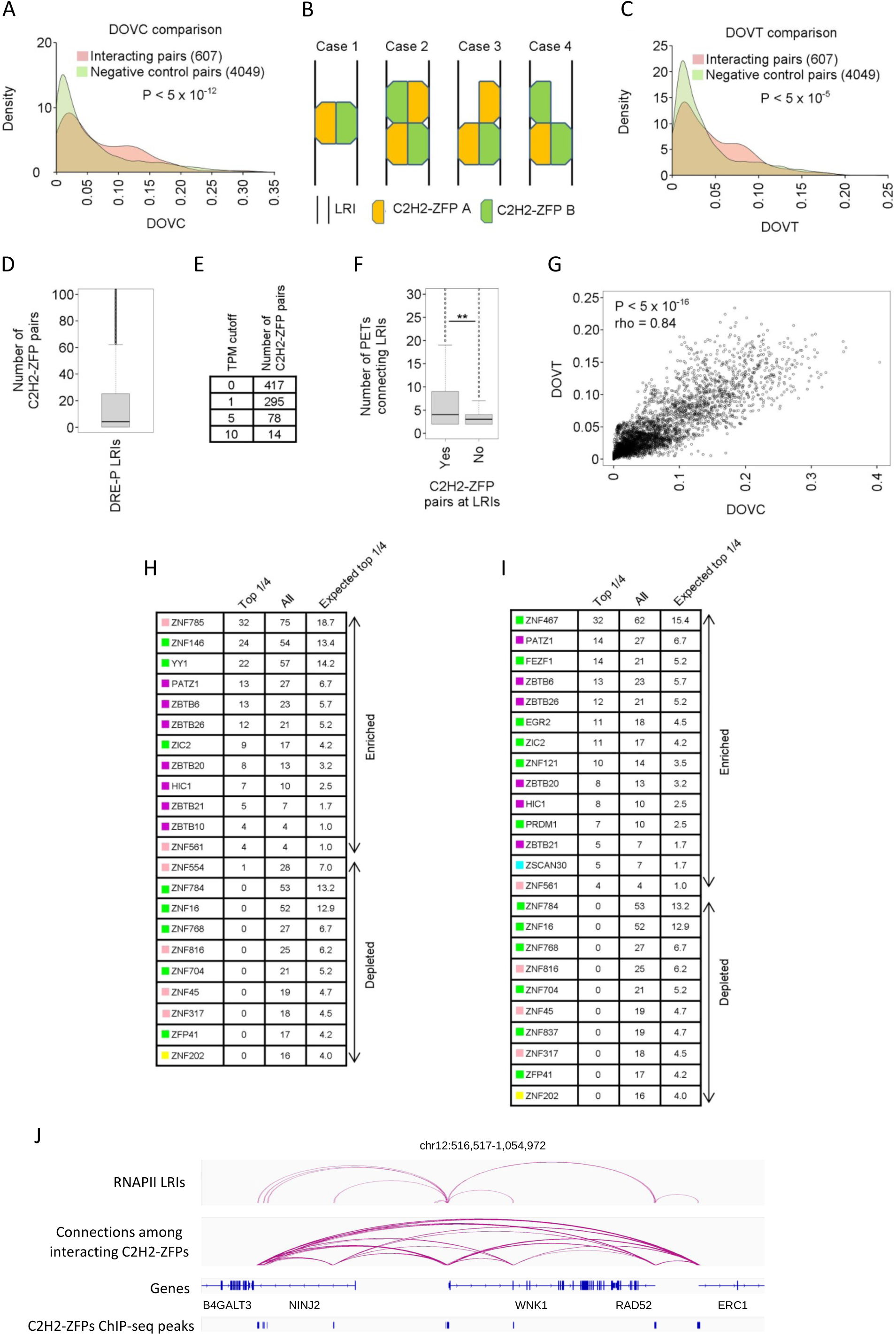
Interacting C2H2-ZFP proteins co-localize significantly at the same as well as at the opposite ends of LRIs. A) Density plots comparing distributions of DOVC values for interacting C2H2-ZFP pairs from LUMIER to negative controls. The DOVC values were calculated using only DNA-binding sites that overlap with LRI anchors. p-values were calculated using the Mann-Whitney test. Negative controls consist of all pairwise C2H2-ZFP combinations that were not found to interact by LUMIER. (B) Scenarios where two C2H2-ZFPs might be found to bind to LRIs. (C) Density plots comparing distributions of DOVT values for interacting C2H2-ZFP pairs to negative controls at LRIs. p-values were calculated using the Mann-Whitney test. Negative controls consist of all pairwise C2H2-ZFP combinations that were not found to interact by LUMIER. (D) Boxplot showing the distribution of the numbers of different C2H2-ZFP pairs that bind DRE-P LRIs on opposite anchors. (E) Table showing the number of extrapolated C2H2-ZFP pairs that may bind at LRIs. The extrapolated numbers have been calculated by using a range of TPM values determined from the median expression of C2H2-ZFPs across tissues. (F) Boxplots comparing the distributions of the number of PETs that link LRIs that have at least one bound C2H2-ZFP pair at opposite LRI anchors (Yes) to LRIs that do not (No). ** p-value < 5 x 10^-16^ (Mann-Whitney test). (G) Plot displaying the correlation of the DOVT and DOVC values for interacting C2H2-ZFP pairs. (H-I) Tables listing significantly enriched or depleted C2H2-ZFPs among the top ¼ pairs compared to the rest according to the ranking by DOVC (H) or DOVT (I) values (p-value < 0.05; hypergeometric test). For each C2H2-ZFP, the table indicates the actual (Top 1/4) and expected (Expected top 1/4) frequency among the top ¼ as well as the frequency among all pairs (All). The colored boxes indicate C2H2-ZFP subfamilies as in Fig. 1B. (J) Genome browser snapshot of selected locus showing the potential connections among interacting C2H2-ZFPs. Top panel: RNAPII LRIs from ChIA-PET dataset. Second panel (from the top): Connections among interacting C2H2-ZFPs that bind at the anchors of the RNAPII LRIs. Third panel: Locations and names of genes in the depicted locus. Fourth panel: Location of the ChIP-seq summits for the interacting C2H2-ZFPs in the second panel.

Second, we evaluated co-binding *in trans* by measuring the frequency with which interacting pairs of C2H2-ZFP occupy opposite anchors of the same chromatin interactions (referred to as ‘degree of overlap *in trans*’ [DOVT]), as detected by ChIP-seq data. For each pair, we quantified the fraction of LRIs bound by both proteins under four possible binding configurations (Fig. 4B; Supplemental Fig. S4D; Supplemental Table S10). These include: 1) protein A and protein B bind at opposite ends of the LRI, 2) both proteins bind at both ends of the LRI, 3) protein A binds at both ends of the LRI but protein B binds at only one end, 4) and *vice versa*. Consistently, interacting pairs exhibit significantly higher DOVT values than an equal number of negative control pairs (Fig. 4C; Supplemental Fig. S4E). Importantly, this enrichment is lost upon randomization of LRI positions (Supplemental Fig. S4F). Finally, utilizing both ChIP-seq and ChIP-exo datasets, AP/MS-derived interacting pairs also exhibit significantly higher DOVT values compared to negative controls (Supplemental Fig. S4G). Together, these results indicate that interacting C2H2-ZFPs preferentially co-occupy chromatin loop anchors, both locally and across interacting genomic loci.

### Multiple interacting C2H2-ZFP pairs converge on individual chromatin interactions

Given the above-described results, we next examined whether chromatin interactions are typically associated with single or multiple C2H2-ZFP pairs. Our analysis revealed that the median number of interacting C2H2-ZFP pairs bound per LRI was four (Fig. 4D). Given that we analyzed only a subset of C2H2-ZFPs and extrapolating based on the number of C2H2-ZFPs expressed in cells (Fig. 4E), we estimate that individual LRIs may be occupied by at least 14 interacting pairs (see methods). This suggests that LRIs are bound by multiple C2H2-ZFP pairs, so these pairs might cooperatively function in LRI formation and/or stabilization. In line with this, we found that LRIs bound by at least one interacting C2H2-ZFP pair had significantly more PETs than LRIs lacking such pairs (Fig. 4F), reinforcing a relationship between interaction strength and occupancy by pairs of interacting C2H2-ZFPs.

We also examined the overall rankings of the interacting pairs based on their DOVC and DOVT values and observed a strong correlation (Fig. 4G). Notably, proteins enriched among high-ranking pairs were predominantly BTB-C2H2-ZFPs, including PATZ1, ZBTB6, ZBTB26, ZBTB20, HIC1 and ZBTB21 (Fig. 4H, I). Figures 4J and S4H depict example loci demonstrating potential connections among the interacting C2H2-ZFPs. Together, these findings suggest that common subsets of C2H2-ZFPs participate in coordinated *cis* and *trans* binding at chromatin loop anchors.

### Binding sites of LRI-associated C2H2-ZFPs are recurrently mutated in cancer

Previous studies have shown that CTCF binding sites at chromatin loop anchors are frequently mutated in cancer (57, 58). Given our above-described results indicating a widespread association of C2H2-ZFPs with LRIs, we next wanted to examine whether the DNA binding sites of such C2H2-ZFPs at loop anchors are similarly affected in cancers. To this end, for each C2H2-ZFP, peak summits within LRI anchors were intersected with somatic mutations from cancer genomes (59) and compared with multiple background models. Specifically, our background controls included: 1) a random chromosome-matched set of locations for the C2H2-ZFP ChIP-seq peak summits within LRI anchors, 2) HEK293 cell DNase I hypersensitive sites within LRI anchors, and 3) HEK293 cell DNase I hypersensitive sites along with the ChIP-seq peaks of all 216 C2H2-ZFPs within LRI anchors (Supplemental Table S1). The overlap analyses were done separately for DRE and promoter anchors. When we employed all above-noted background strategies, we found 51 and 45 C2H2-ZFPs whose DNA binding sites significantly overlap cancer mutations at distal and promoter anchors, respectively (Fig. 5A, Supplemental Table S11). In total, DNA-binding sites for 75 of 216 C2H2-ZFPs (∼35%) show significant overlap with cancer mutations at either anchor type. This set included CTCF, as well as many poorly characterized C2H2-ZFPs (Fig. 5A-C), several of which have few prior links to cancer biology (Fig. 5B).

**Figure 5.**
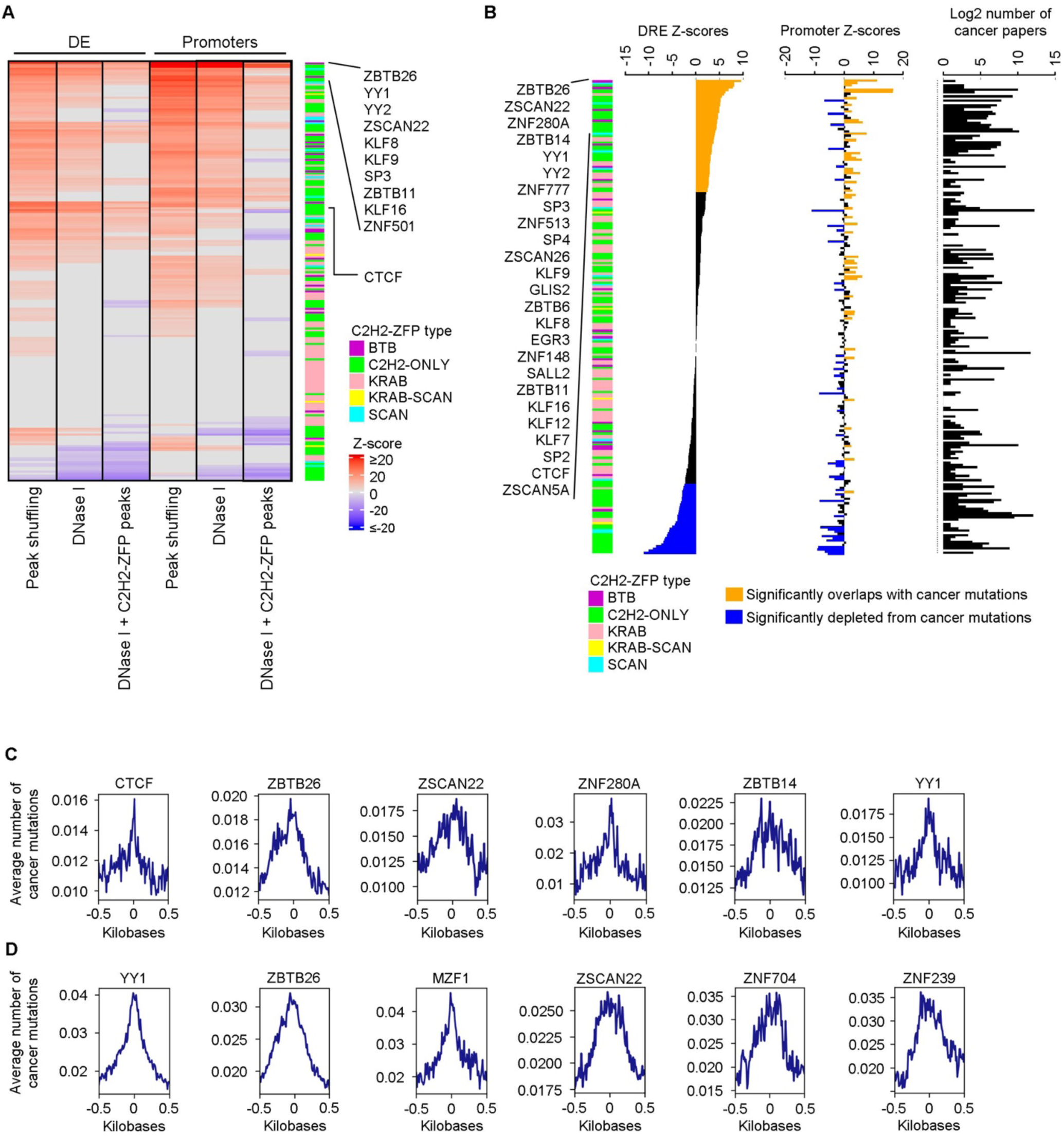
Binding sites of C2H2-ZFPs at LRI anchors are mutated in cancer. (A) Heatmap demonstrating Z-scores for the overlap of DNA-binding of C2H2-ZFPs with cancer mutations across DRE and promoter LRI anchors. The overlaps were done using three different null backgrounds for the C2H2-ZFP ChIP-seq datasets, which consisted of 1) a random chromosome-matched set of locations for the C2H2-ZFP ChIP-seq peaks within LRI anchors (Peak shuffling), 2) HEK293 DNase I hypersensitive sites within LRI anchors (DNase I), and 3) HEK293 DNase I hypersensitive sites along with the ChIP-seq peaks of all 216 C2H2-ZFPs analyzed in this study within LRI anchors (DNase I + C2H2-ZFP peaks). Only C2H2-ZFPs with significant Z-scores are displayed. (B) Barplots displaying Z-scores for the overlap of the DNA-binding of C2H2-ZFPs with cancer mutations across DRE and promoter LRI anchors, as well as the number of papers that have been published for the C2H2-ZFPs related to cancer. The displayed Z-scores are the ones that were calculated using the DNase I + C2H2-ZFP peaks background in panel A. (C-D) Metaplot representations of cancer mutations around ChIP-seq peak summits for some selected C2H2-ZFPs at (C) DRE and (D) promoter LRI anchors +/- 500 bp. These examples were selected to display cases where mutations were more concentrated close to the centers of the peak summits and cases where the cancer mutation were more dispersed around the peak summits.

We next performed mutation distribution analysis and observed distinct patterns across various C2H2-ZFPs. For example, for CTCF, ZNF280A and YY1, mutations were concentrated near peak summits, whereas for ZBTB26, ZSCAN22 and ZBTB14 they were more broadly distributed (Fig. 5C; Supplemental Fig. S5). Similar patterns were observed at promoter anchors, e.g., for YY1 and MZF1 the cancer mutations were more concentrated around the peak summits whereas for ZSCAN22 and ZNF704 they were more dispersed (Fig. 5D; Supplemental Fig. S6). The dispersed mutation profiles suggest that, in some cases, mutations may affect not only the binding sites of individual C2H2-ZFPs but also those of their interacting partners. Together, these results suggest that a substantial fraction of LRI-associated C2H2-ZFP binding sites are recurrently mutated in cancer genomes, implicating these factors, and perhaps their interaction networks, in disease-associated regulatory disruption.

## Discussion

Although C2H2-ZFPs comprise the largest class of human TFs, the majority of the family members and their functional relevance have remained poorly characterized (4). In this study, we report the protein-protein interaction network for more than half of all human C2H2-ZFPs, finding that these factors interact extensively with one another, in addition to their interactions with proteins involved in diverse functions, ranging from transcriptional cofactors, RNA-binding proteins, and chromatin-associated factors to proteins involved in mitochondrial homeostasis. We further show that a substantial fraction of the C2H2-ZFPs that we analyzed localize to anchors of LRIs and that their binding at these sites is correlated with changes in the expression of associated genes. Our study also revealed that interacting pairs of C2H2-ZFPs are enriched both at the same loop anchors and at opposite ends of chromatin loops, raising the interesting possibility that C2H2-ZFPs might function in the formation or stability of LRIs through homodimerization and heterodimerization. Finally, we show that the binding sites of many LRI-associated C2H2-ZFPs are recurrently mutated in cancer, implicating disruption of these regulatory interactions in transcriptional dysregulation.

By using complementary approaches, we identified hundreds to thousands of pairwise interactions among C2H2-ZFPs. For example, AP/MS detected 278 high-confidence interactions, whereas LUMIER uncovered a substantially broader pairwise interaction landscape. Because C2H2-ZFPs tend to be expressed at relatively low levels compared with many other nuclear proteins in HEK293 cells (Supplemental Fig. S7), it is likely that AP/MS underestimated interactions that involve low-abundance factors, as suggested previously for proteomics studies (56). In contrast, LUMIER assays involve overexpression of both proteins in cells, which may increase sensitivity but potentially at the cost of introducing non-physiological associations. Nevertheless, the key conclusion from these analyses, i.e., that C2H2-ZFPs extensively interact with each other, is supported by both methods.

An intriguing question that arises from the extensive connectivity observed among C2H2-ZFPs relates to the nature of the complexes that these proteins may form. Neither AP/MS nor LUMIER resolves whether interactions are direct or mediated through higher-order assemblies, nor do they provide information about stoichiometry. While some C2H2-ZFPs are known to self-associate, it is also plausible that distinct family members form heteromeric or oligomeric structures. Similar assemblies have been described for factors such as SP1 and PRDM9 (24, 60). Thus, it is imaginable that related mechanisms may operate more broadly within this family. Structural and biochemical studies will be required to define the interaction interfaces and organization of these complexes. Another potential implication of our findings is that the presence of multiple C2H2-ZFP pairs at LRI anchors may facilitate the formation of higher-order assemblies through weak, multivalent protein-protein and protein-DNA interactions. Such interaction networks have been proposed to underlie phase separation or condensate-like organization at regulatory regions of the genome (61, 62). In this scenario, C2H2-ZFPs and their associated cofactors could contribute to the establishment of locally enriched regulatory compartments at LRIs that support chromatin looping and gene regulation. Testing this model will require future studies that directly probe the biophysical properties of C2H2-ZFP assemblies *in vivo*.

Chromatin loops play an important role in gene regulation, as they can bring regulatory elements, e.g., enhancers or promoters, physically close to their target genes (19, 63). Among the C2H2-ZFP family, CTCF was shown to be a critical factor that brings distant genomic elements into close spatial proximity by driving chromatin loop formation (43, 47). The ability of CTCF to drive chromatin looping has been shown to depend on dimerization of two CTCF molecules that are bound in convergent orientation on distant genomic elements (64). In this regard, our study suggests that several additional C2H2-ZFPs likely function in chromatin looping. This is supported by our observation that interacting pairs of C2H2-ZFPs frequently bind at opposite ends of LRIs. Moreover, we found that multiple distinct interacting pairs frequently converge on the same chromatin interaction. Although our estimates are derived from bulk data and likely overstate the number of factors bound simultaneously at individual LRIs in single cells, they nevertheless suggest that chromatin loops can be occupied by coordinated assemblies of C2H2-ZFPs. The correlation between co-binding in *cis* and co-occupancy in *trans* further supports a model in which C2H2-ZFP interactions contribute to loop architecture. It should be noted, however, that the DNA binding profiles in our analyses were derived from ChIP-seq experiments that were done using ectopically expressed proteins. This may introduce non-physiological occupancy, particularly in accessible chromatin. To mitigate this, we focused on reproducible peaks and employed multiple background controls, yet we cannot fully rule out residual bias. Another limitation is that we used ChIP-seq peak summits as proxies for binding sites; while these are typically enriched for cognate motifs (65), they do not guarantee direct DNA contact. Previous studies have suggested that PPIs among TFs can enable recognition of alternative DNA-binding motifs (66). Given the extensive PPIs among C2H2-ZFPs, it is plausible to assume that at least some genomic associations might have been caused by indirect recruitment through interacting partners rather than direct motif recognition. Future studies should aim at dissecting the role of interacting C2H2-ZFP pairs in chromatin looping by using more direct approaches. For example, it will be important to perform Hi-C or similar assays in cells where LRI-associated C2H2-ZFPs are depleted to examine their effect on LRIs.

Moreover, C2H2-ZFPs can also associate with RNA, and several recent studies have shown that many human C2H2-ZFPs directly bind RNA *in vivo* (67–70). This raises the possibility that RNA-mediated interactions could further influence the chromatin association and regulatory activity of C2H2-ZFPs. This idea is consistent with previous findings indicating that RNA binding by CTCF is important for its role in LRIs (71, 72). How RNA binding by the C2H2-ZFPs that we have identified here intersects with their DNA binding and protein-protein interactions at LRIs remains an open question and may represent an additional layer of regulation.

Chromatin loops have been implicated in number of gene expression regulatory processes (73). For example, recent studies show that, in addition to impacting the transcription of target genes, CTCF-mediated chromatin loops might also regulate alternative splicing (74, 75). Consistent with a functional role at chromatin loops, we found that depletion of several C2H2-ZFPs altered the expression of genes associated with the LRIs they occupy. The direction and magnitude of these effects varied among factors and depended in some cases on whether binding occurred at promoters, distal regulatory elements, or both. These findings are consistent with context-dependent regulatory roles, and a number of scenarios can be envisioned; for example, C2H2-ZFPs may promote loop formation between promoters and enhancers to activate transcription or may restrict interactions between promoters and distal elements to insulate or repress gene expression. Furthermore, differential recruitment of cofactors in distinct genomic contexts is also likely to contribute to these opposing outcomes. Further studies are needed to investigate these various possibilities.

Finally, we show that binding sites of many LRI-associated C2H2-ZFPs significantly overlap somatic mutations in cancer genomes. This extends previous observations for CTCF (58) and suggests that disruption of C2H2-ZFP-mediated regulatory interactions may be a recurrent feature of tumor genomes. Such mutations may impair loop formation or stability and thereby alter gene regulatory programs. These findings are consistent with a growing body of evidence suggesting that disease-causing DNA mutations can alter the ability of TFs to recognize their cognate binding sites (76, 77). The set of C2H2-ZFPs identified here as loop-associated and mutation-enriched provides a framework for future studies aimed at defining how perturbation of chromatin architecture contributes to oncogenic transcriptional dysregulation. In this regard, it will be interesting to explore how often do C2H2-ZFPs’ binding sites overlap common variants without known pathogenicity? And whether C2H2-ZFP overlap common variants more than the binding sites of other TFs do?

Based on our results, we suggest a model in which C2H2-ZFPs form interconnected interaction networks that assemble at chromatin loop anchors (Fig. 6). Such PPIs could bridge (or stabilize) distal regulatory elements (e.g., enhancers and promoters), which may contribute to the establishment or maintenance of LRIs. Furthermore, we suggest that, depending on the genomic context and cofactors that might be recruited, C2H2-ZFPs may either promote or restrict regulatory communication between distal elements and target genes. Disruption of these interactions by somatic mutation may therefore compromise chromatin architecture and gene regulation in cancer.

**Figure 6.**
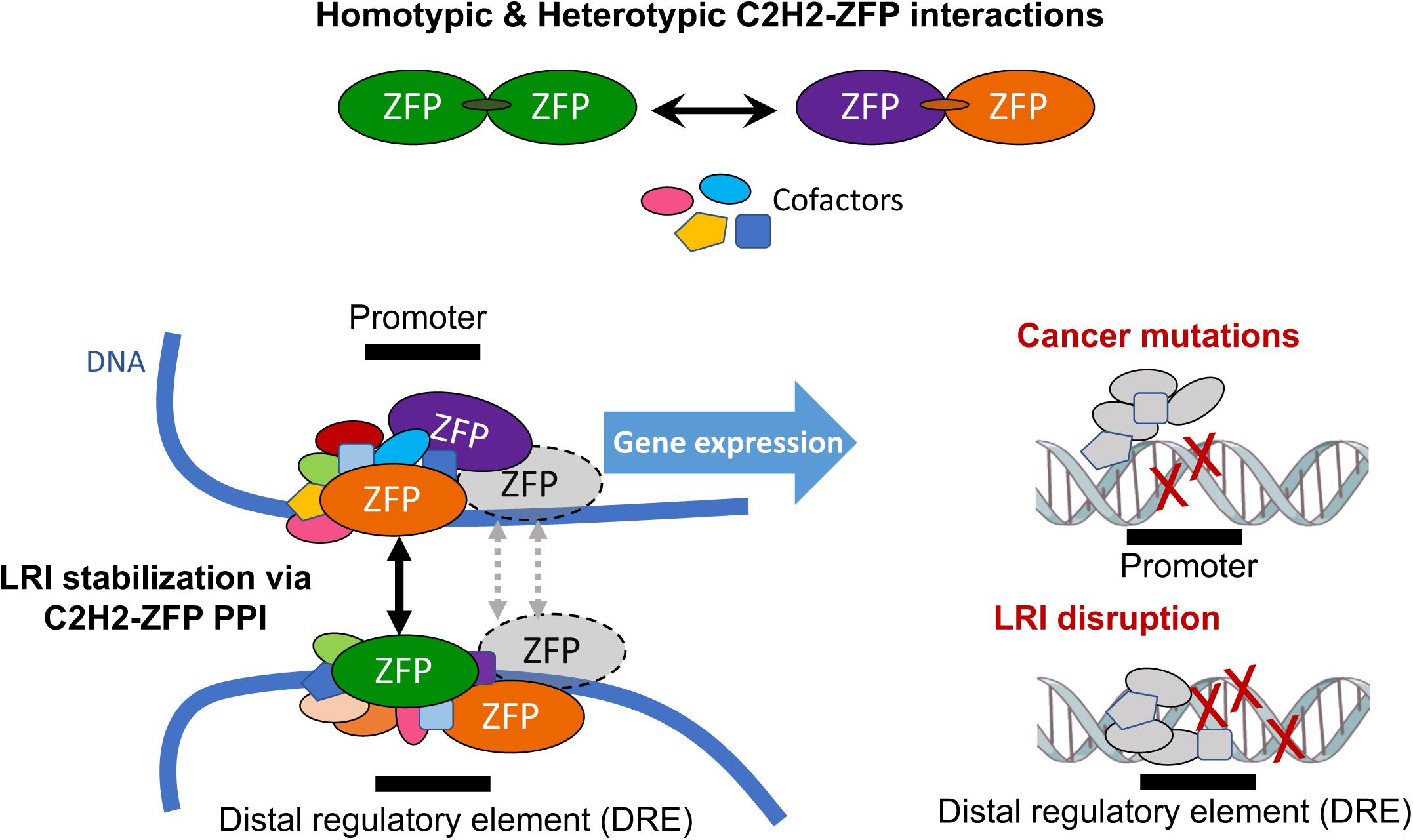
Model for co-operative C2H2-ZFP function at long-range chromatin interactions (LRIs). C2H2-ZFPs bind at both promoters and distal regulatory elements (DREs) that participate in LRIs. Extensive protein–protein interactions among C2H2-ZFPs enable their co-binding at individual LRI anchors and, in some cases, at opposite ends of the same loop. These interactions may contribute to the formation or stabilization of LRIs and thereby influence gene expression. Somatic mutations that overlap C2H2-ZFP binding sites at LRI anchors may perturb these interactions and compromise transcriptional regulation. Note: C2H2-ZFPs shaded in grey, as well as grey dotted lines, indicate that multiple C2H2-ZFPs may participate in LRI regulation. Moreover, proteins depicted in grey in the cancer mutation panel highlight that certain co-factors (or non-C2H2-ZFP transcription factors) might be recruited to these sites despite the absence of C2H2-ZFPs.

## Material and Methods

### Annotation of human promoters

The hg38 GENCODE release 44 .gtf annotation file (77) was used as a reference for construction of promoters of protein-coding genes. All promoters consist of the region from 1,000 bp upstream of the transcription start site to 500 bp downstream of the transcription start site of the longest isoforms of protein-coding genes. Protein-coding genes are cases where in the .gtf annotation file, column 3 is labeled as “gene” and column 12 is labeled as “protein_coding”.

### Identification of RNAP II bound long range interactions

Replicate paired end unmapped reads for RNAP II in situ ChIA-PET datasets (ENCLB238DBN and ENCLB885AAO) were obtained from the ENCODE consortium portal (Supplemental Table S1). ChIA-PET Tool V3 (44) was used to identify LRIs from the paired end raw reads for each biological replicate. The ChIA-PET Tool V3 was run in long-read mode with default settings. The forward and reverse bridge linker sequences provided to the program were ACGCGATATCTTATCTGACT and AGTCAGATAAGATATCGCGT, respectively, and the option --minimum_linker_alignment_score 14 was used as recommended by the authors (44). Following the identification of significant LRIs from each biological replicate (adjusted p-value < 0.05), inter-chromosomal LRIs were removed and only the reproducible ones were selected for downstream analysis. This was done by using bedtools pairtopair (78) to select only LRIs identified from ENCLB885AAO that had both anchors overlap with both anchors of LRIs from ENCLB238DBN. The LRIs were then classified into 1) promoter-DRE, 2) promoter-promoter, and 3) DRE-DRE. Promoter-DRE LRIs were ones where one of the anchors had at least one base pair overlapping with a promoter of a protein-coding gene and the other anchor did not. Promoter-promoter LRIs were ones where both LRI anchors had at least one base pair overlapping with promoters of protein-coding genes. DRE-DRE LRIs were ones where neither anchor had any overlap with promoters of protein coding genes.

### DNase I hypersensitivity data processing

DNase I hypersensitivity data was processed by mapping reads, filtering reads, calling peaks, and evaluating which peaks were reproducible. DNase I data from HEK293T cells consisting of six biological replicates was downloaded from Gene Expression Omnibus entry GSE67007 (79, 80). cutadapt version 1.9.1 was used to trim reads to 20 bp (81, 82). Next, reads were mapped to the hg38 genome assembly and filtered using the ATAC-Seq / DNase-Seq Pipeline (https://github.com/kundajelab/atac_dnase_pipelines) with options - dnase_seq, -align, and -multimapping 4 (83). Since all datasets seemed high-quality, biological replicates were merged into two larger pooled replicates containing three replicates each such that the pooled replicates were as close in size as possible. Peaks were then called, reproducible peaks were identified, and signal tracks from mapped reads were created using the ATAC-Seq / DNase-Seq Pipeline with option -enable_idr. Reproducible peaks (IDR < 0.1) across pooled replicates were used for downstream analyses (84).

### Functional element overlap

To determine the significance of the overlap of a C2H2-ZFP with a genomic element at the DNA level with the peak shuffling background, the expected overlap of the C2H2-ZFP with the genomic element was calculated by shuffling the ChIP-seq peak summits 1,000 times in the same chromosomes as their original locations and ensuring that shuffled summits do not overlap with each other using bedtools shuffle (version 2.29.0) with the -chrom - noOverlapping options (78). After each shuffle, the overlap with the functional element was calculated. A normal distribution was fitted to these overlaps, which was then used to obtain a mean and standard deviation that was used to calculate the empirically derived Z-score for the actual number of summits that overlapped the genomic element for each C2H2-ZFP. The overlaps were identified using bedtools intersect (version 2.29.0). For the overlaps using as a null background HEK293 DNase I hypersensitive sites or HEK293 DNase I hypersensitive sites along with the ChIP-seq peaks of all 216 C2H2-ZFPs analyzed in this study, instead of randomly shuffling the C2H2-ZFP ChIP-seq peak summits across the genome, peak summit sets the same size as the peak sets were randomly drawn for each C2H2-ZFP from these two collections of peaks. For all the overlaps, the p-values were calculated using the pnorm function in R, and the results were corrected for multiple hypothesis testing with the Benjamini-Hochberg method by using the p.adjust function in R with the option “BH”. Heatmaps displaying the Z-scores from the overlaps were generated with the R package ComplexHeatmap (85).

### RNA-seq data analysis

Publicly available FASTQ files for datasets where the expression of C2H2-ZFPs had been disrupted as well as associated controls were obtained from sources listed in Supplemental Table S1. RSEM (86). Reads were mapped using STAR (87) with the option --star, and default parameters were used to generate gene level quantification files. The genome and transcriptome from GENCODE release 46 was used as a reference. The counts from the gene level quantification files were used to estimate the changes in gene expression between treatment and control experiments using DESeq2 (88). Significantly mis-regulated genes were considered ones that had an adjusted p-value from DESeq2 < 0.05.

### Generation of cell lines

C2H2-ZFP ORFs (open reading frames) were cloned into the pDEST pcDNA5/FRT/TO eGFP vector through Gateway™ cloning (Invitrogen) using procedures recommended by the manufacturer. Successfully tagged genes were verified by restriction enzyme digestion of the vector. A list of the origins of the constructs that were used in this study can be found in Supplemental Table S3. Flp-IN TREx HEK293 cells (Invitrogen) were used to generate cell lines expressing GFP tagged proteins as previously described (14).

### AP/MS method

Single-step affinity purifications of GFP-tagged proteins in HEK293 cells were done as previously described (14). Briefly, ∼20 million cells were harvested following a 24h induction of protein expression by adding doxycycline to the media. After preparing whole-cell extracts, they were incubated overnight with an anti-GFP antibody (G10362, Life Technologies), followed by a two-hour incubation with Protein G Dynabeads (Invitrogen) to immunoprecipitate GFP-tagged proteins. Following washes, the immunoprecipitated material was eluted with ammonium hydroxide and digested with trypsin. The samples were then desalted and analyzed with an LTQ-Orbitrap Velos mass spectrometer (ThermoFisher Scientific).

### SAINT analysis of PPI data

The supervised scoring approach, SAINTexpress, which compares putative PPIs to a negative control set of purifications (48), was used to assign confidence values to the PPIs. This scoring system takes as input total spectral counts of prey proteins and assigns each prey protein a probability of interaction. All the AP/MS interactions used in this study were derived from an analysis where we used as negative controls 28 AP/MS experiments done on GFP, 171 experiments done on 78 different non-C2H2-ZFP TFs from various subfamilies, and 17 AP/MS experiments done on 9 different chromatin-related proteins (Supplemental Fig. S2B and Supplemental Table S1). However, below, we also compared this analysis to one where we used as background only 28 AP/MS experiments done on GFP, because conventionally, purifications done with only the tag are used as a background in analyses scoring PPIs from AP/MS. To quantitatively assess the performance of the SAINT score resulting from the use of different backgrounds, a Gold Standard reference set of PPIs from the BioGRID database (89, 90) was used. For an interaction to be included in the Gold Standard reference set, it had to be supported by at least one high-throughput and one low-throughput experiment. The only high-throughput experiment taken into consideration was AP/MS, whereas low-throughput experiments taken into consideration consisted of the following BioGRID tags: (1) Affinity Capture followed by Western blotting, (2) Co-Crystal Structure, (3) Reconstituted Complex, (4) Co-purification, (5) Affinity Capture-Luminescence, and (6) Protein Peptide. To construct the negative control set, random pairs between bait and prey proteins were selected such that they do not correspond to any known human PPIs in BioGRID. The size of the negative control set was 10 times larger than that of the positive control set, which consisted of 416 Gold Standard reference PPIs among 124 unique baits and 279 unique preys.

When GFP purifications (GFP-ONLY) were used as negative controls, even at the most stringent SAINT cutoff of 1, the precision achieved, which is defined as (True positives)/(True positives + False positives), was only 0.787 (Supplemental Fig. S8A). This low precision likely occurred because GFP fails to fully capture the background, especially the chromatin-related one, since it is not a DNA-binding protein. For example, proteins such as MDC1, NUMA1, PRMT5, TOP2A, WDR77, XRCC6, and others tend to appear in high spectral counts in C2H2-ZFP purifications, but very low counts in GFP purifications (Supplemental Fig. S8D), and, after SAINT scoring with GFP-ONLY as a negative control set, at cutoffs of 0.93 and 1, most interactions with these proteins score as significant (Supplemental Fig. S8E). In the past, to deal with this issue, such proteins that frequently score as significant across purifications have been labelled as “frequent fliers” and have been manually removed from analyses (14). However, many of these proteins have key roles in chromatin dynamics, so eliminating them from analyses may result in the lack of detection of important interactions that impact gene expression. These same proteins also appear to be present at similar levels across 188 purifications for 78 non-C2H2-ZFP TFs and 9 chromatin-related proteins (Supplemental Fig. S8E), which were purified in parallel with the C2H2-ZFPs using the same method. When the non-C2H2-ZFP TFs were added to the negative control set, along with the GFP purifications (GFP + TF), most interactions with the frequent fliers were not significant at a SAINT cutoff of 1 or 0.93 (Supplemental Fig. S8E). At a SAINT cutoff of 0.93, interactions with frequent fliers that scored as significant involved cases where the frequent fliers were highly enriched compared to other C2H2-ZFP purifications (Supplemental Fig. S8F).

ROC (receiver operator characteristic) curve analysis shows that SAINT with GFP + TFs as background, at a cutoff of 0.93, has a better FPR (FPR = 0.00120) than when using GFP-ONLY as background (FPR = 0.00829) (Supplemental Fig. S8B). While at a SAINT cutoff of 0.93 the use of the GFP-ONLY background results in better recall, it also results in worse precision compared to the GFP + TF background. A precision of 1 can only be achieved if the SAINT score cutoff used is 0.99 in the analysis with GFP + TFs as background, which also reduces the recall to 0.0817 (Supplemental Fig. S8A). Although using GFP + TFs as background detects fewer true positives than when using GFP-ONLY as background, it provides a much better precision. Therefore, to achieve a good balance between precision and recall in our analysis the GFP+TF background at a SAINT score cutoff of 0.93 was used to determine the significant interactions (FDR = 0.109) (Supplemental Fig. S8C).

### Filtering of potentially misidentified AP/MS peptides

To minimize any potential artifacts in the mass spectrometry data for C2H2-ZFP interactions, the following peptides were removed from the analysis: 1) incompletely digested peptides 2) peptides that had a difference of less than or equal to two amino acids from the best matching sequence in the bait, not including a difference between leucine and isoleucine, and 3) peptides that had any detected posttranslational modifications. The homology of prey peptides with the bait protein sequence was assessed using ClustalW. Furthermore, to minimize interactions due to carryover, prey C2H2-ZFPs were removed from the network if they were also analyzed as baits in the same series of MS experiments where they showed up as prey. A summary of the filtered PPIs can be found in Supplemental Figure S3A, and a full list of removed peptides and interactions can be found in Supplemental Table S8.

### DAVID analysis and cluster generation

The analyses with DAVID were carried out by using default parameters on the DAVID online portal (53) on August 29, 2023. The cutoff used to select significant functional terms was a Bonferroni corrected p-value < 0.05. The terms used to describe the selected clusters in Supplemental Tables S5 and S6 were chosen to broadly reflect the terms significantly enriched for each cluster. The age of C2H2-ZFPs was determined based on the analysis done in Figure S3A by Lambert et al. (1). Great Apes C2H2-ZFPs were defined as ones that were found in at least one great ape (Human, Chimpanzee, or Gorilla) but were found in 30% or fewer of the other species in the figure. Primate C2H2-ZFPs were defined as ones that were found in at least one primate (Human, Chimpanzee, Gorilla, Macaque, Marmoset, or Mouse Lemur) but were found in 30% or fewer of the other species in the figure. Old C2H2-ZFPs were defined as ones that were found in over 70% of non-primate species in the figure. A C2H2-ZFP was considered to be found in a species if it had a non-zero % amino acid identity. The identity of C2H2-ZFPs belonging to each age group can be found in Supplemental Table S13.

### LUMIER experimental procedure

The LUMIER assay protocol was employed as described previously (55). Expression plasmids for bait proteins and prey proteins, viral packaging and generation of stable cell lines were all done as described (55). HEK293 cell lines were used for all experiments. Stable cell lines expressing prey proteins were seeded into 96-well plates. When the cells reached 80% confluency (30,000 cells) they were transfected with 150 ng of a plasmid expressing 3xFLAG-tagged bait protein or a negative control plasmid without FLAG using PEI transfection reagent. LUMIER assay plates coated with FLAG M2 antibody were prepared on the day of transfection. After 24 hours, cells were lysed by addition of AFC buffer (10 mM Tris HCl, pH 7.9, 150 mM NaCl, 0.1% NP40, protease inhibitors, 10mM NaF, and 2 mM sodium pyrophosphate) for 10 minutes, incubating on ice. Plates were then sonicated 2×10 min using an in-bath sonicator (VWR, model 750) at high power. After sonication, samples were incubated for 1 hour on ice and 60 ml of cell lysate was then transferred to the assay plate coated with anti-FLAG antibody. The plates were incubated for 3 hours at 4°C and washed with 250 ml of cold AFC buffer, 4 times, with 5 min incubation time each time. All the remaining steps were carried out as described in Taipale (2018) on pages 53-54.

### LUMIER data analysis

1. Data scaling: First, LUMIER intensity values were log-transformed. Two different negative controls were used, H2O and cells expressing Flag-tagged GFP. These negative controls were used to control for non-specific binding and to account for batch-to-batch effects. Subsequently, the mean and standard deviation of data points corresponding to the negative controls were computed. All data points were subsequently scaled using these mean and standard deviations to obtain LUMIER scores:

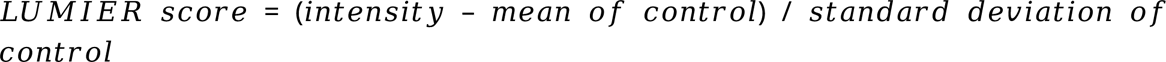

2. Cut-off determination: The smallest LUMIER score that produced a 15% false discovery rate (FDR, equivalent to an 85% precision) was chosen as the cut-off. Specifically, LUMIER scores of positive controls on multiple plates corresponding to known interactions (between TRIM28 and ZNF669) were used as the positive reference set. The negative reference set matching the size of the positive reference set was randomly chosen from LUMIER scores of all samples in a treatment group, based on the assumption that true interactions are rare, and the vast majority of the LUMIER scores represent background noise associated with non-specific binding. At a given LUMIER score, the positive reference set would be partitioned into true positives (TP, those above the score) and false negatives (FN, those below the score); meanwhile, the negative reference set would similarly be partitioned into false positives (FP) and true negatives (TN). The false discovery rate is calculated as the following:

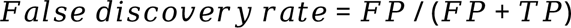

LUMIER scores were tested in an ascending order by calculating their associated FDRs, and the first score that produced an FDR less than 15% was considered as a cut-off. To avoid sampling bias, 1,000 negative reference sets in total were generated by random sampling, and a cut-off was determined for each of them. The median of the resulting 1,000 cut-offs was selected as the final cut-off for the entire dataset.

Following the analysis of the raw LUMIER data, the modifications described below were done to remove non-C2H2-ZFP proteins from the analysis, deal with multiple names for the same protein and combine PPIs obtained from multiple constructs from the same protein. ILF2, TRIM21, TRIM28, ZFY1, ZRANB2, ZSWIM2_C2, ZSWIM2_C1, ZZZ3_C2 were removed from the final C2H2-ZFP PPI list because they are not C2H2-ZFPs. EGR2_C1, EGR2_C2,KLF6_C1, KLF6_C2,PRDM1_C1, PRDM1_C2, PRDM1_C3, REPIN1_C1, REPIN1_C2, REPIN1_C3, REPIN1_C4, SP2_C1, SP2_C2, ZBTB10_C1, ZBTB10_C2, ZBTB18_C1, ZBTB18_C2, ZBTB20_C1, ZBTB20_C2, ZBTB44_C1, ZBTB44_C2, ZFP28_C1, ZFP28_C2, ZNF146_C1, ZNF146_C2, ZNF195_C1, ZNF195_C2, ZNF273_C3, ZNF273_C4, ZNF3_C1, ZNF3_C3, ZNF34_C1, ZNF34_C2, ZNF423_C1, ZNF423_C2, ZNF432_C1, ZNF432_C2, ZNF512_C1, ZNF512_C2, ZNF529_C1, ZNF529_C2, ZNF544_C1, ZNF544_C2, ZNF554_C1, ZNF554_C2, ZNF71_C1, ZNF71_C2, ZSCAN16_C1, ZSCAN16_C2, ZSCAN20_C1 and ZSCAN20_C2 were cases where multiple constructs were used to test PPIs for the same gene (“C” is an abbreviation for construct); therefore, the PPIs were combined together by gene name in the final interaction list (Supplemental Table S9B). Because C2H2-ZFPs could have more than one name, in two cases two different constructs with different names for the same proteins were used in the assay. Those two cases were dealt as follows: 1) ZBTB14 appears in bait proteins as ZBTB14 and ZFP161; therefore, ZFP161 was replaced with ZBTB14 in baits, and duplicate interactions were removed in the final PPI list (Supplemental Table S9B). The name ZFP161 was changed to ZBTB14 also in preys. 2) ZNF192 was changed to ZKSCAN8 in the prey list because ZNF192 was used as the name when the TF was a prey, whereas ZKSCAN8 was used as the name when the TF was a bait. Furthermore, to match more up-to-date names used for two C2H2-ZFPs, ZNF816A was changed to ZNF816 and ZNF397OS was changed to ZSCAN30 in Supplemental Table S9B. Homodimers and reciprocal interactions were removed from the analysis with LRI overlaps, but they were kept for the heatmap generation in Fig. 3D. When interactions were identified in duplicate, the one with the highest Z-score was used to generate the heatmap.

### Overlap of AP/MS and LUMIER data

The hypergeometric test was used to determine whether AP/MS confirmed more C2H2-ZFP/C2H2-ZFP LUMIER interactions than expected by chance, and *vice versa*. To determine whether AP/MS confirmed more C2H2-ZFP/C2H2-ZFP LUMIER interactions than expected by chance, the expected number of interactions for the 345 bait proteins was calculated by assuming expression of prey proteins at nTPM cuttoffs of 0, 1 and 5 while keeping the number of the identified PPIs constant. The nTPM values for C2H2-ZFPs in HEK293 cells were obtained from the Human Protein Atlas (91). For both analyses, only unique interactions were considered, and homodimeric interactions were removed from the analysis since AP/MS cannot identify them.

### ChIP-seq data analysis

The ENCODE IDR optimal C2H2-ZFP ChIP-seq peaks were obtained from the ENCODE portal (Supplemental Table S1) (45). The dataset described here also includes existing eGFP-ChIP-seq data from HEK293 Flp-IN T-REx cells, which were obtained from the Gene Expression Omnibus (80, 92) entries GSE52523 (13) and GSE76496 (14). These data are single-end instead of paired-end. These data were reprocessed in the same way as the ENCODE data using the AQUAS Transcription Factor and Histone ChIP-Seq processing pipeline (https://github.com/kundajelab/chipseq_pipeline) with the hg38 genome assembly and default settings for single-end reads, which filters out multi-mapping reads (93). The control was a pool of all the input datasets with Normalized Strand Cross-correlation coefficient (NSC) < 1.05 after running the AQUAS pipeline through the cross-correlation step (93–95), randomly subsampled to fifty million reads. For several C2H2-ZFPs, one replicate’s raw reads file had been corrupted (IKZF3, ZNF16, ZNF213, ZNF524, ZNF543, ZNF594, ZNF610, ZNF677, ZNF98, ZSCAN30, and ZSCAN5C); for these, the reads from the bam files of reads that mapped to hg19 were extracted, where bam files were obtained from the authors of Schmitges et al. (14), and the pipeline was run starting with those reads. The fourth replicate of SP4 from Schmitges et al. (14)was also excluded because both the raw reads and bam files had been corrupted. In addition, for datasets with more than two biological replicates, between the read filtering and peak-calling steps, all biological replicates with NSC < 1.05 were removed; if more than two replicates remained, the remaining replicates were combined into two groups of “pooled replicates” such that the groups had the same number of filtered reads. If a dataset had only one biological replicate, either initially or after removing low-quality replicates, the reads were divided randomly into two “pseudo-replicates,” and the IDR reproducible peaks across the pseudo-replicates were obtained. For all other datasets, the “IDR optimal set” was used (84).

### ChIP-exo data processing

ChIP-exo peaks for 28 C2H2-ZFPs (Supplemental Table S1) were obtained from Imbeault et al. (15). For ZNF100, ZNF419 and ZNF282, which had datasets from multiple biological replicate experiments, the ChIP-exo peaks from the different replicates were combined by using bedtools merge. The liftOver tool (96, 97) was then used to convert coordinates for each dataset from hg19 to hg38 so that they could be overlapped with the other datasets that were mapped to hg38. For the analyses involving calculation of DOVC values, the midpoint of the ChIP-exo peaks was used as a summit to perform the 150bp extensions for generating the peaks used in the analyses. For the analyses involving the calculation of the DOVT values, the ChIP-exo peaks were used as provided without performing any extensions.

### Calculation of DOVC values

Co-binding for two hypothetical proteins A and B at LRI anchors was quantified with the DOVC value, which was calculated with the following formula:

DOVC = (Number of ChIP-seq peaks at LRI anchors overlapping between protein A and B)/((Total Number of ChIP-seq peaks at LRI anchors for protein A) + (Total Number of ChIP-seq peaks at LRI anchors for protein B)).

For the C2H2-ZFP pairs from LUMIER, the DOVC values were calculated after extending the C2H2-ZFP ChIP-seq peak summits 150 bp on each side. The overlaps were identified by using bedtools intersect (78), and any degree of overlap between two ChIP-seq peaks was considered to be an overlap. p-values that assessed the difference in distributions of DOVC values between interacting C2H2-ZFPs and ones that were not found to interact were calculated by using the Mann-Whitney test in R. Interacting C2H2-ZFPs consisted of 607 pairs formed by 97 unique C2H2-ZFPs for which ChIP-seq data was publicly available (13, 14, 46). Negative control C2H2-ZFP pairs consisted of all the pairwise combinations of the 97 C2H2-ZFPs that were not found to interact with each other based on LUMIER.

The analyses for the C2H2-ZFP pairs from AP/MS were performed similarly to the analyses described for the LUMIER pairs. In these analyses, interacting C2H2-ZFPs consisted of 94 pairs formed by 85 unique C2H2-ZFPs, for which ChIP-seq or ChIP-exo data was publicly available (13–15, 46). Negative control C2H2-ZFP pairs consisted of all the pairwise combinations of the 85 C2H2-ZFPs that were not found to interact with each other based on AP/MS.

### Calculation of DOVT values

To quantify the overlap of C2H2-ZFPs with LRIs for two given C2H2-ZFPs, A and B, the DOVT value was calculated using the formula:

DOVT = (LRIs bound by A and B)/((LRIs bound by A) + (LRIs bound by B) - (LRIs bound by A and B)).

An LRI was considered to be bound by a pair of C2H2-ZFPs if (1) the two C2H2-ZFPs bound only at opposite ends, (2) one of the C2H2-ZFPs bound both ends, whereas the other bound only one end, or (3) both C2H2-ZFPs bound both ends. An LRI was considered to be bound if there was any overlap between the LRI anchors and C2H2-ZFP ChIP-seq peak summits. p-values, which assessed the difference in distributions of DOVT values between interacting C2H2-ZFPs and ones that were not found to interact, were calculated using the Mann-Whitney test in R. Interacting C2H2-ZFP pairs from LUMIER consisted of 607 pairs formed by 97 unique C2H2-ZFPs, where ChIP-seq data was publicly available (13, 14, 46) for both interacting proteins in the pairs. Negative control C2H2-ZFP pairs consisted of all the pairwise combinations of the 97 C2H2-ZFPs that were not found to interact with each other based on the LUMIER data. The same procedure was followed to overlap the interacting C2H2-ZFP pairs, as well as the negative controls, with 1,000 sets of shuffled LRIs. The DOVT values from all the overlaps with the shuffled LRIs were then combined for the two sets of C2H2-ZFP pairs, and their distributions were compared to the distribution of the DOVT values generated from the overlap of interacting pairs with the actual LRIs by using the Mann-Whitney test. Density plots and boxplots showing the distributions were generated using R. This same procedure was used to overlap AP/MS pairs with LRIs, with the only difference that ChIP-exo experiments from Imbeault et al. (15) (Supplemental Table S1) were added to the analysis so that ChIP-seq data was available for 94 pairs involving 85 C2H2-ZFPs. Negative control C2H2-ZFP pairs consisted of all the pairwise combinations among the 85 C2H2-ZFPs that were not found to interact with each other based on AP/MS.

### Identification of C2H2-ZFPs that are significantly enriched and depleted among pairs that colocalize most significantly *in cis* and *in trans*

To determine the significantly enriched and depleted C2H2-ZFPs among the pairs that colocalize most significantly *in cis* in Figure 4H, the pairs were first ranked based on their DOVC values, and the top ¼ of the pairs were considered as the ones that colocalize most significantly *in cis*. The p-values were calculated by using the hypergeometric test in R, and they were corrected using the Benjamini-Hochberg method. A similar procedure was followed to determine the C2H2-ZFPs that were significantly enriched and depleted among pairs that colocalize most significantly *in trans*, with the only difference that the pairs were ranked based on the DOVT values from the overlaps with the LRIs indicated in Figure 4C.

### Calculation of the number of C2H2-ZFP pairs at LRIs

For each C2H2-ZFP pair, the unique LRIs that were bound on opposite ends were identified by using bedtools pairtobed. An LRI was considered to be bound by a pair of proteins if (1) the two proteins bound only at opposite ends, (2) one of the proteins bound both ends whereas the other bound only one end, or (3) both proteins bound both ends. For these overlaps, the ChIP-seq peak summits of C2H2-ZFPs were used. After the overlaps with each individual pair, the bound LRIs were combined and the frequency with which a particular LRI appeared in the combined set was considered as the number of C2H2-ZFP pairs that were bound at the LRI.

### Overlap of C2H2-ZFP DNA-binding with cancer mutations

The cancer mutations for the overlaps with C2H2-ZFP binding sites were obtained from COSMIC (cancer.sanger.ac.uk) and they were from the version 101 of their files (59). The set of mutations at DRE anchors was constructed from the hg38 version of the file containing the non-coding variants. If the mutation appeared in the file it was only counted once, and any overlapping mutations were merged using bedtools merge (78). For each C2H2-ZFP, peak summits that were extended 15bp on each side were overlapped with mutations at DREs as described for the overlaps with functional elements with the only difference that the shuffling of the peaks and the selection of random DNase I sites as well as C2H2-ZFP peak summits was done within the boundaries of the DRE anchors. Similarly, to the DRE anchor mutations, the set of mutations at promoter anchors was constructed from the hg38 version of the file containing the non-coding variants, but the coding mutations from the Census Genes Mutations hg38 version of the file were also included. Census Genes Mutations were included in this case because, based on our definition, promoters extend 500bp downstream of the TSSs, which is a region that can contain coding elements of genes. The overlaps with C2H2-ZFP DNA-binding sites were done similarly to the ones with the DRE anchor cancer mutations with the only difference that the overlaps were restricted to those with anchors that overlapped with promoters. Metaplots showing the distribution of cancer mutations relative to C2H2-ZFPs peak summits were constructed with deeptools (98).

### Pubmed searches

Pubmed searches were done on January 21, 2025. To identify the number of cancer-related papers for every C2H2-ZFP, the gene name along with the phrase “and cancer” was searched in Pubmed. The initial search for REST yielded an excessive number of publications. Because the name of this C2H2-ZFP is also a verb and a noun, most papers were not relevant, and the alternative name of the gene, NRSF, was used instead.

Processing of RNA-seq data for extrapolation of the number of C2H2-ZFP pairs at LRIs

RNA-seq data was obtained from the Human Protein Atlas (https://www.proteinatlas.org/) on August 7, 2023 (91). The normalized TPM values that were provided were used to calculate the median expression for each gene across tissues. Data was used from all tissues, except for the amygdala, basal ganglia, cerebellum, hippocampal formation, hypothalamus, midbrain, pituitary gland, spinal cord, and vagina, because data from these tissues was provided for fewer genes (19,266 versus 20,162).

Extrapolation of the number of C2H2-ZFP pairs at LRIs was performed as following: Our LUMIER experiments tested 54 bait proteins against 283 prey proteins, yielding 1,732 unique interactions from 15,282 pairwise tests. Assuming a similar interaction frequency across the C2H2-ZFP family, we estimated the potential number of interactions among expressed C2H2-ZFPs. For example, among the 137 C2H2-ZFPs expressed at TPM >10, this interaction frequency would predict approximately 2,127 pairwise interactions. In our integrative analysis we examined 607 interacting pairs and observed four pairs binding opposite ends of DRE–promoter LRIs. Extrapolating from the estimated interaction space suggests that as many as ∼14 pairs of C2H2-ZFPs could potentially occupy opposite ends of LRIs. Similar calculations performed using broader expression thresholds (TPM >5, >1, or all C2H2-ZFPs) yielded estimates ranging up to several hundred potential pairs at LRIs.

### Comparison of expression of C2H2-ZFP to non-C2H2-ZFP proteins

The quantification of protein expression by mass spectrometry was obtained from Geiger et al. (99). The Log10 intensity score after iBAQ normalization was used as a measure for protein expression. Given that experiments were done in triplicate, the average Log10 intensity score was calculated for each protein. If a quantification score was obtained in only one of the replicate experiments, then that value was used as the protein expression level. The Mann-Whitney test was then used to compare the distributions of the protein expression levels, and the boxplots were generated with R.

## Acknowledgments

We thank Frank Schmitges, Lixia Jiang-Xu and Anshul Kundaje for useful discussions about ChIP-seq. We thank Abraham Weintraub, Denesz Hnisz and Richard Young for guidance through ChIA-PET. We thank Jeff Liu for data storage support and Peter Young for experimental support. Team members at the Donnelly Sequencing centre are gratefully acknowledged for their help with next generation sequencing.

## Authors’ contribution

ER and JFG conceived the study. ER participated in cell line generation and AP/MS, performed all omics’ data analysis, and contributed to manuscript drafting. EM and HT participated in LUMIER assays. S.N-S wrote the manuscript, participated in data analysis, and co-ordinated edits from all authors. SP participated in MS and LUMIER data analysis. GZ participated in cell line generation, tissue culture, and AP/MS. HG participated in MS experiments. IMK contributed to data analysis. IMK, AE and TRH took part in manuscript editing. JFG supervised the project and edited the manuscript.

## Data access

All raw AP/MS datasets generated in this study have been submitted to the MassIVE repository. All the raw and processed LUMIER datasets generated in this study have been provided as supplementary material with the paper.

