## Supplemental figures for "Integrative analysis reveals extensive interactions among C2H2 zinc finger proteins at chromatin loop anchors"

### Supplemental Figure 1

A

| Statistic being reported | Replica 1 | Replica 2 |
|---|---|---|
| Total PETs | 529,138,548 | 339,527,099 |
| Valid PETs after linker filtering | 382,075,028 | 234,529,648 |
| Uniquely Mapped same-linker PETs | 316,591,088 | 197,367,605 |
| Merging same same-linker PETs | 72,994,659 | 41,220,546 |
| Merging similar same-linker PETs | 61,475,616 | 35,913,991 |
| Self-ligation PETs | 28,182,358 | 14,690,011 |
| Inter-ligation PETs | 12,909,604 | 7,532,434 |
| Other PETs with short distance | 20,383,654 | 13,691,546 |
| Peaks from self-ligation PETs | 5,815 | 4,439 |
| Signficant long range interactions | 729,107 | 624,133 |
| Signficant interchromosomal long range interactions | 109,733 | 78,528 |
| Signficant intrachromosomal long range interactions | 619,374 | 545,605 |
| Reproducible singificant long range interactions | 63,511 | |
| Significant interchromosomal long range interactions that are reproducible | 177 | |
| Significant intrachromosomal long range interactions that are reproducible | 63,334 | |

B

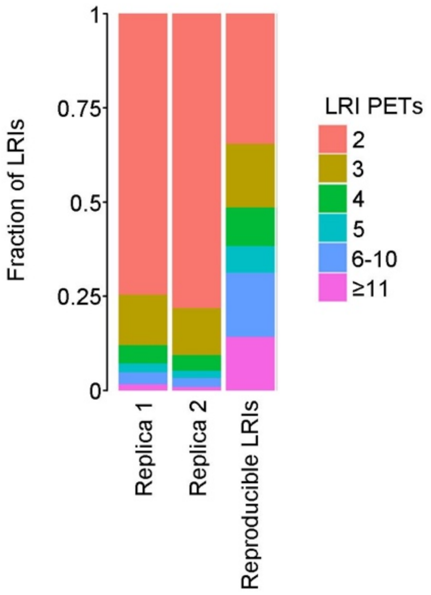


C

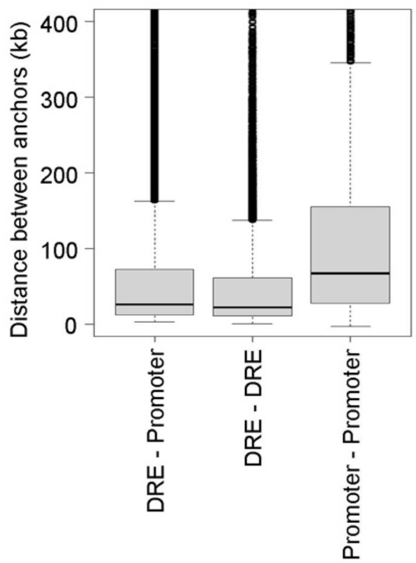


**Supplemental Figure 1. Analysis of RNAP II ChIA-PET data.**
(A) Table listing some statistics on the analysis of the two RNAP II ChIA-PET datasets in HEK293 cells.
(B) Barplot showing the fraction of LRIs that are linked by the indicated number of PETs for the reproducible LRIs, as well as the LRIs that were identified from the two individual RNAP II ChIA-PET replicate experiments.
(C) Boxplots showing the distribution of the distance between two different anchors for the indicated LRI types

### Supplemental Figure 2

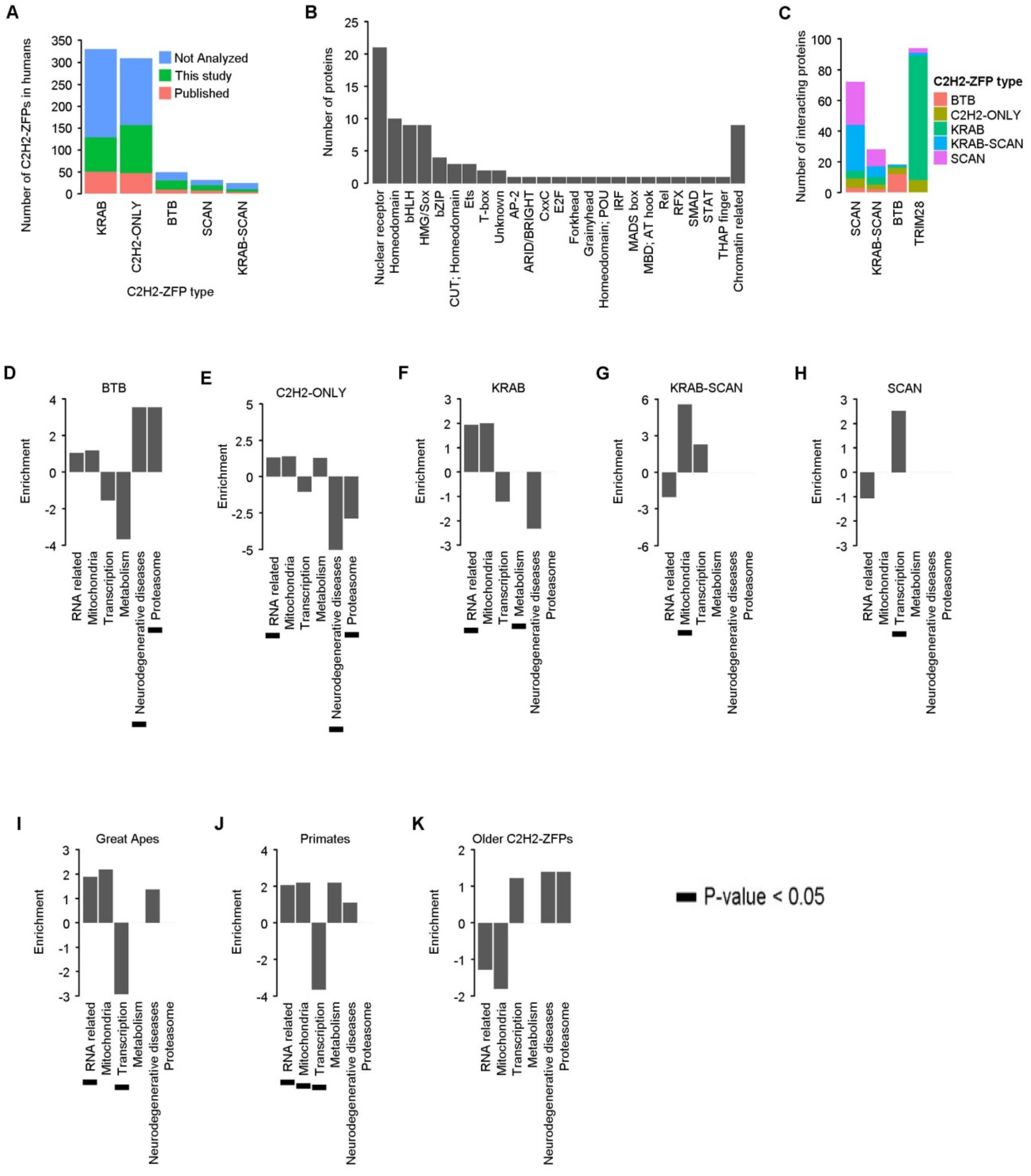

**Supplemental Figure S2. PPI network of C2H2-ZFPs uncovers diverse interactions involved in multiple biological processes.**

**(**A) Summary of C2H2-ZFP subfamilies that were characterized by AP/MS.

(B) Barplot showing the subfamilies to which the TFs used as background in the SAINT analysis belong. The 9 non-TF proteins used in the background were classified as "Chromatin related".

(C) Barplot showing that AP/MS for C2H2-ZFPs captures many PPIs with other C2H2-ZFPs that are expected for C2H2-ZFPs based on auxiliary domains. For TRIM28 as a prey protein, or for baits containing a KRAB, SCAN or BTB domain, the types of interacting C2H2-ZFPs are classified according to their auxiliary domains.

(D-K) Barplots showing the enrichment of selected functional terms among PPI preys of (D) BTB-C2H2-ZFPs, (E) C2H2-ONLY-ZFPs, (F) KRAB-C2H2-ZFPs, (G) KRAB-SCAN-C2H2-ZFPs, (H) SCAN-C2H2-ZFPs, (I,J) C2H2-ZFPs found predominantly in (I) Great Apes or (J) Primates and (K) older C2H2-ZFPs. The p-values were calculated using the hypergeometric test, and they were corrected using the Benjamini-Hochberg method.

### Supplemental Figure 3

**A**

| SAINT score | Good preys | Possible peptide misidentifications | Possible carryovers | Percentage of filtered preys |
|---|---|---|---|---|
| 1 | 101 | 14 | 15 | 22.3 |
| 0.99 | 51 | 11 | 1 | 19.1 |
| 0.98 | 10 | 1 | 0 | 9.1 |
| 0.97 | 52 | 22 | 4 | 33.3 |
| 0.96 | 11 | 2 | 0 | 15.4 |
| 0.95 | 4 | 0 | 1 | 20.0 |
| 0.94 | 41 | 41 | 0 | 50.0 |
| 0.93 | 9 | 1 | 0 | 10.0 |
| Total | 279 | 92 | 21 | 28.8 |

**B**

| | C2H2-ONLY (35) | KRAB (43) | KRAB-SCAN (6) | SCAN (12) | BTB (8) |
|---|---|---|---|---|---|
| C2H2-ONLY (27) | 36 | 11 | 2 | 2 | 3 |
| KRAB (41) | 7 | 117 | 0 | 2 | 1 |
| KRAB-SCAN (8) | 3 | 8 | 7 | 30 | 1 |
| SCAN (6) | 0 | 1 | 11 | 25 | 0 |
| BTB (10) | 3 | 0 | 0 | 1 | 8 |

**C**

| | C2H2-ONLY (61) | KRAB (89) | KRAB-SCAN (13) | SCAN (13) | BTB (26) |
|---|---|---|---|---|---|
| C2H2-ONLY (18) | 254 | 368 | 69 | 50 | 98 |
| KRAB (20) | 249 | 334 | 70 | 57 | 96 |
| KRAB-SCAN (1) | 16 | 15 | 8 | 8 | 6 |
| SCAN (1) | 5 | 4 | 4 | 4 | 0 |
| BTB (5) | 56 | 49 | 11 | 7 | 27 |

**D**

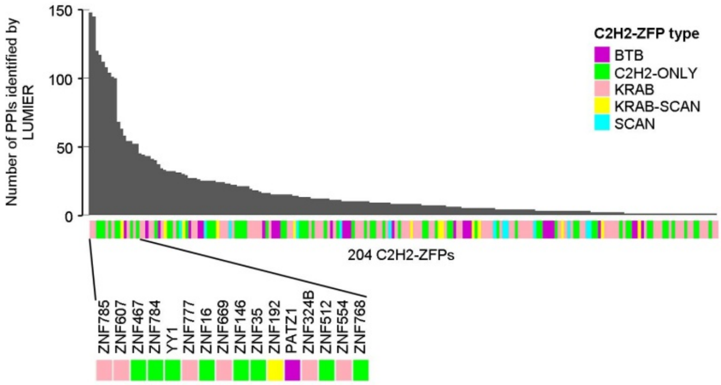


**E**

| | |
|---|---|
| Bait proteins | 282 |
| Prey proteins | 54 |
| Proteins tested as both bait and prey | 52 |
| Interactions tested by LUMIER | 13850 |
| Interactions confirmed by LUMIER | 1732 |
| AP-MS PPIs tested by LUMIER | 74 |
| AP-MS PPIs confirmed by LUMIER | 21 |
| AP-MS PPIs expected to be confirmed by chance | 9.25 |
| P-value | <0.0005 |

**F**

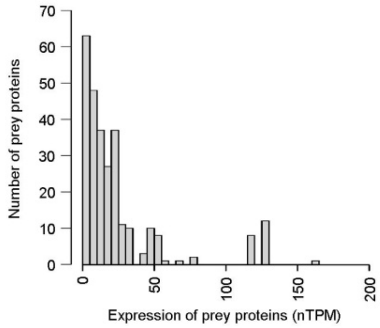


**G**

| TPM cutoff for prey proteins | 0 | 1 | 5 |
|---|---|---|---|
| Bait proteins | 345 | 345 | 345 |
| Prey proteins | 751 | 609 | 401 |
| Interactions tested by AP/MS | 199410 | 150420 | 78660 |
| Interactions confirmed by AP/MS | 278 | 278 | 278 |
| LUMIER interactions tested by AP/MS | 1656 | 1656 | 1656 |
| LUMIER interactions confirmed by AP/MS | 21 | 21 | 21 |
| LUMIER interactions expected to be confirmed | 2.31 | 3.06 | 5.85 |
| p-value | $<5\times10^{-14}$ | $<5\times10^{-11}$ | $<5\times10^{-7}$ |

**Supplemental Figure S3. C2H2-ZFPs interact with each other extensively.**

(A) Table summarizing the filtering of PPIs due to prey peptides that could have been misidentified by mass spectrometry at different SAINT cutoffs.

(B) Summary of the number of PPIs among C2H2-ZFPs from different subfamilies based on the AP/MS network from Fig. 3C. The numbers inside brackets in the column headings represent the number of unique bait C2H2-ZFPs displaying PPIs. The numbers inside brackets in the row labels represent the number of unique prey C2H2-ZFPs identified as interactors.

(C) Summary of the number of PPIs among C2H2-ZFPs from different subfamilies based on LUMIER PPIs from Fig. 3D. The numbers inside brackets represent the same types of C2H2-ZFPs as in panel B.

(D) Barplot displaying the number of interactions that LUMIER identified for each C2H2-ZFP that was tested.

(E) Table with statistics assessing whether LUMIER confirms more AP/MS C2H2-ZFP PPIs than expected by chance. The p-value was calculated using the hypergeometric test.

(F) Histogram displaying the distribution of expression levels of prey C2H2-ZFP proteins in HEK293 cells that appear in the C2H2-ZFP interacting pairs that were identified by AP/MS. C2H2-ZFPs that are members of pairs that could have been misidentified because of mass spectrometry artifacts are not included.

(G) Table with statistics assessing whether AP/MS confirms more LUMIER C2H2-ZFP PPIs than expected by chance. Calculations were done by using different TPM values as a cutoff for the expression of prey C2H2-ZFPs in HEK293 cells. The p-values were calculated using the hypergeometric test .

### Supplemental Figure 4

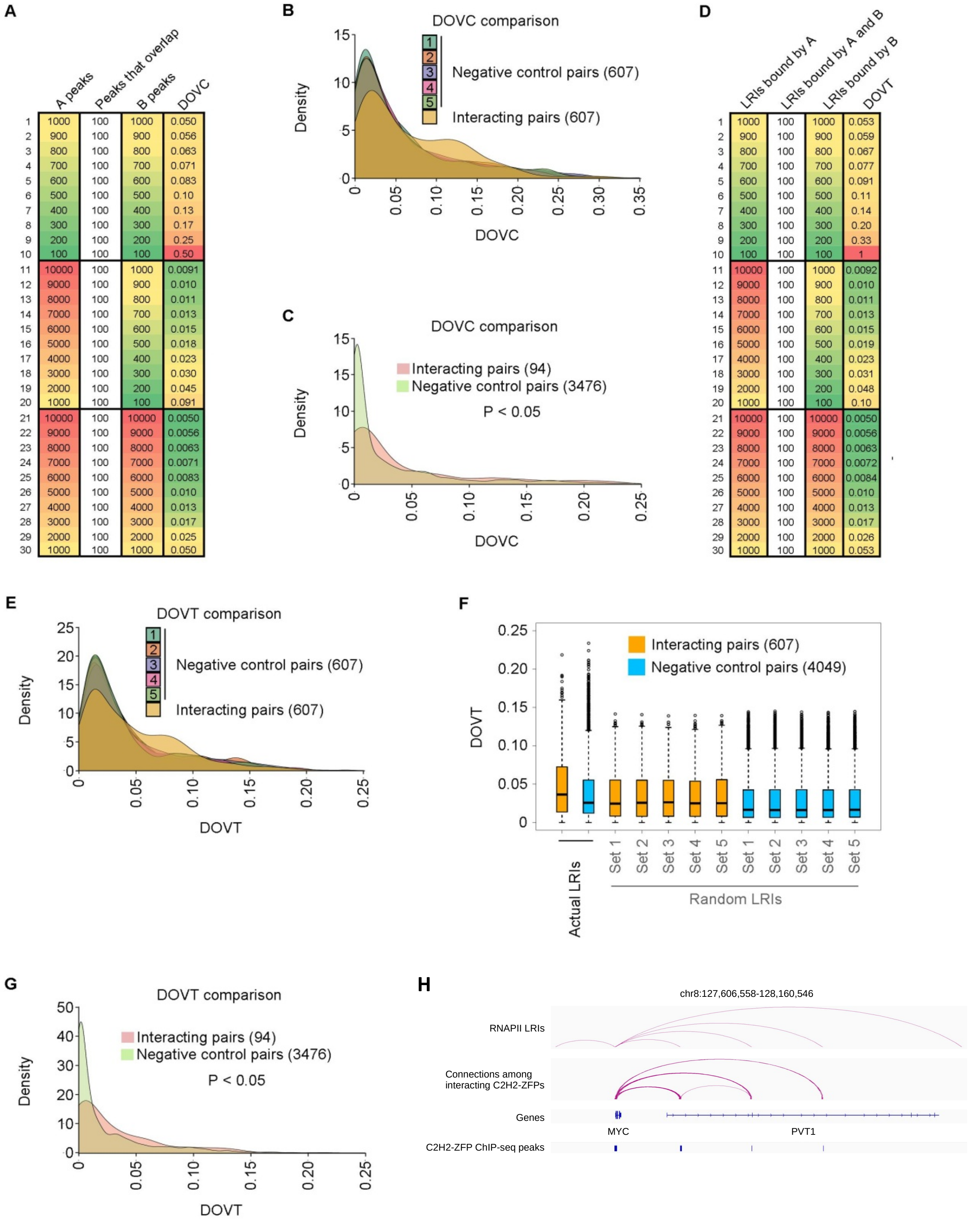


**A**

| | A peaks | Peaks that overlap | B peaks | DOVC |
|---|---|---|---|---|
| 1 | 1000 | 100 | 1000 | 0.050 |
| 2 | 900 | 100 | 900 | 0.056 |
| 3 | 800 | 100 | 800 | 0.063 |
| 4 | 700 | 100 | 700 | 0.071 |
| 5 | 600 | 100 | 600 | 0.083 |
| 6 | 500 | 100 | 500 | 0.10 |
| 7 | 400 | 100 | 400 | 0.13 |
| 8 | 300 | 100 | 300 | 0.17 |
| 9 | 200 | 100 | 200 | 0.25 |
| 10 | 100 | 100 | 100 | 0.50 |
| 11 | 10000 | 100 | 1000 | 0.0091 |
| 12 | 9000 | 100 | 900 | 0.010 |
| 13 | 8000 | 100 | 800 | 0.011 |
| 14 | 7000 | 100 | 700 | 0.013 |
| 15 | 6000 | 100 | 600 | 0.015 |
| 16 | 5000 | 100 | 500 | 0.018 |
| 17 | 4000 | 100 | 400 | 0.023 |
| 18 | 3000 | 100 | 300 | 0.030 |
| 19 | 2000 | 100 | 200 | 0.045 |
| 20 | 1000 | 100 | 100 | 0.091 |
| 21 | 10000 | 100 | 10000 | 0.0050 |
| 22 | 9000 | 100 | 9000 | 0.0056 |
| 23 | 8000 | 100 | 8000 | 0.0063 |
| 24 | 7000 | 100 | 7000 | 0.0071 |
| 25 | 6000 | 100 | 6000 | 0.0083 |
| 26 | 5000 | 100 | 5000 | 0.010 |
| 27 | 4000 | 100 | 4000 | 0.013 |
| 28 | 3000 | 100 | 3000 | 0.017 |
| 29 | 2000 | 100 | 2000 | 0.025 |
| 30 | 1000 | 100 | 1000 | 0.050 |

**D**

| | LRIs bound by A | LRIs bound by A and B | LRIs bound by B | DOVT |
|---|---|---|---|---|
| 1 | 1000 | 100 | 1000 | 0.053 |
| 2 | 900 | 100 | 900 | 0.059 |
| 3 | 800 | 100 | 800 | 0.067 |
| 4 | 700 | 100 | 700 | 0.077 |
| 5 | 600 | 100 | 600 | 0.091 |
| 6 | 500 | 100 | 500 | 0.11 |
| 7 | 400 | 100 | 400 | 0.14 |
| 8 | 300 | 100 | 300 | 0.20 |
| 9 | 200 | 100 | 200 | 0.33 |
| 10 | 100 | 100 | 100 | 1 |
| 11 | 10000 | 100 | 1000 | 0.0092 |
| 12 | 9000 | 100 | 900 | 0.010 |
| 13 | 8000 | 100 | 800 | 0.011 |
| 14 | 7000 | 100 | 700 | 0.013 |
| 15 | 6000 | 100 | 600 | 0.015 |
| 16 | 5000 | 100 | 500 | 0.019 |
| 17 | 4000 | 100 | 400 | 0.023 |
| 18 | 3000 | 100 | 300 | 0.031 |
| 19 | 2000 | 100 | 200 | 0.048 |
| 20 | 1000 | 100 | 100 | 0.10 |
| 21 | 10000 | 100 | 10000 | 0.0050 |
| 22 | 9000 | 100 | 9000 | 0.0056 |
| 23 | 8000 | 100 | 8000 | 0.0063 |
| 24 | 7000 | 100 | 7000 | 0.0072 |
| 25 | 6000 | 100 | 6000 | 0.0084 |
| 26 | 5000 | 100 | 5000 | 0.010 |
| 27 | 4000 | 100 | 4000 | 0.013 |
| 28 | 3000 | 100 | 3000 | 0.017 |
| 29 | 2000 | 100 | 2000 | 0.026 |
| 30 | 1000 | 100 | 1000 | 0.053 |

**Supplemental Figure S4. Interacting C2H2-ZFP proteins co-localize significantly at the same as well as at the opposite ends of LRIs.**

(A) Table showing hypothetical examples of how the DOVC value could change for C2H2-ZFPs when they exhibit different degrees of overlap in ChIP-seq peaks.

(B) Density plots comparing distributions of DOVC values for interacting C2H2-ZFP pairs from LUMIER to negative controls. The DOVC values were calculated using only DNA-binding sites that overlap with LRI anchors. Negative controls consist of five sets of 607 randomly chosen pairwise C2H2-ZFP combinations that were not found to interact by LUMIER. p-values comparing interacting pairs to the five sets of negative controls were calculated using the Mann-Whitney test, corrected using the Benjamini-Hochberg method, and they are listed in Supplemental Table S12A.

(C) Density plots comparing distributions of DOVC values for interacting C2H2-ZFP pairs from AP/MS to negative controls. The DOVC values were calculated using only DNA-binding sites that overlap with LRI anchors. p-values were calculated using the Mann-Whitney test. Negative controls consist of all pairwise C2H2-ZFP combinations that were not found to interact by AP/MS.

(D) Table showing hypothetical examples of how the DOVT value could change for C2H2-ZFPs when they exhibit different degrees of overlap in DNA-binding sites at LRIs.

(E) Density plots comparing distributions of DOVT values for interacting C2H2-ZFP pairs to negative controls. Negative controls consist of five sets of 607 randomly chosen pairwise C2H2-ZFP combinations that were not found to interact by LUMIER. p-values comparing interacting pairs to the five sets of negative controls were calculated using the Mann-Whitney test, corrected using the Benjamini-Hochberg method, and they are listed in Supplemental Table S12B.

(F) Boxplots comparing DOVT values of all interacting C2H2-ZFPs pairs that were identified by LUMIER to various negative controls that include 1) DOVT values that were calculated for all the pairwise combinations of C2H2-ZFPs that were not found to interact by LUMIER using the DRE-promoter LRIs (Actual LRIs), 2) DOVT values that were calculated for all interacting C2H2-ZFP pairs using five sets of shuffled DRE-promoter LRIs, and 3) DOVT values that were calculated for all the pairwise combinations of C2H2-ZFPs that were not found to interact by LUMIER using the same five sets of shuffled DRE-promoter LRIs that were used for the interacting pairs. p-values comparing interacting pairs to negative controls were calculated using the Mann-Whitney test, corrected using the Benjamini-Hochberg method, and they are listed in Supplemental Table S12C.

(G) Density plots comparing distributions of DOVT values for interacting C2H2-ZFP pairs from AP/MS to negative controls at LRIs anchors. p-values were calculated using the Mann-Whitney test. Negative controls consist of all pairwise C2H2-ZFP combinations that were not found to interact by AP/MS.

(H) Genome browser snapshot of selected locus showing the potential connections among interacting C2H2-ZFPs. Top panel: RNAPII LRIs from ChIA-PET dataset. Second panel (from the top): Connections among interacting C2H2-ZFPs that bind at the anchors of the RNAPII LRIs. Third panel: Locations and names of genes in the depicted locus. Fourth panel: Location of the ChIP-seq summits for the interacting C2H2-ZFPs in the second panel.

### Supplemental Figure 5

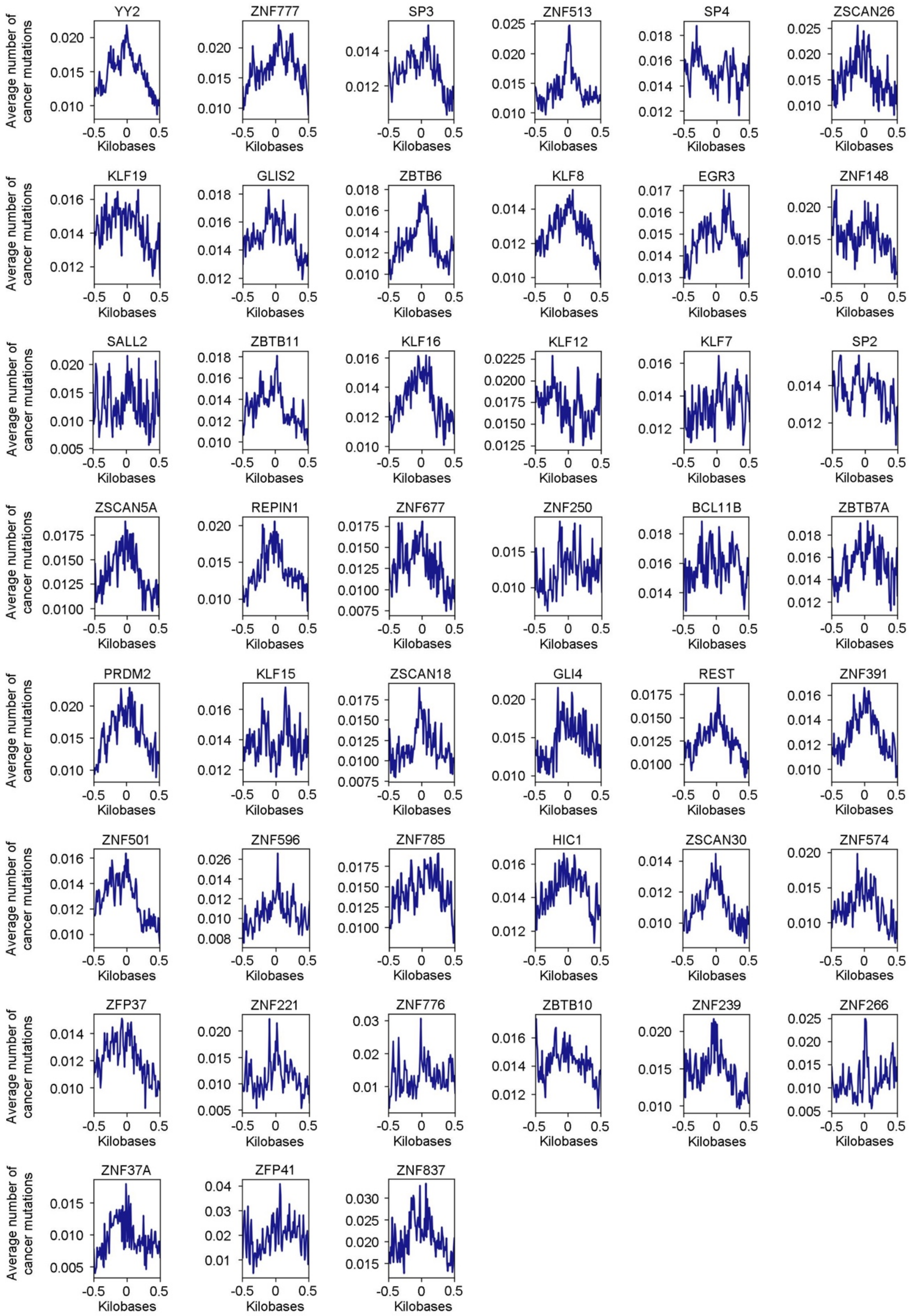

**Supplemental Figure S5. Metaplot representation of cancer mutations around DNA-binding sites for C2H2-ZFPs at DRE LRI anchors +/- 500 bp**. The figure only displays C2H2-ZFPs that overlap significantly with cancer mutations with all three different backgrounds used in Fig. 5A and that were not included in Fig. 5C.

### Supplemental Figure 6

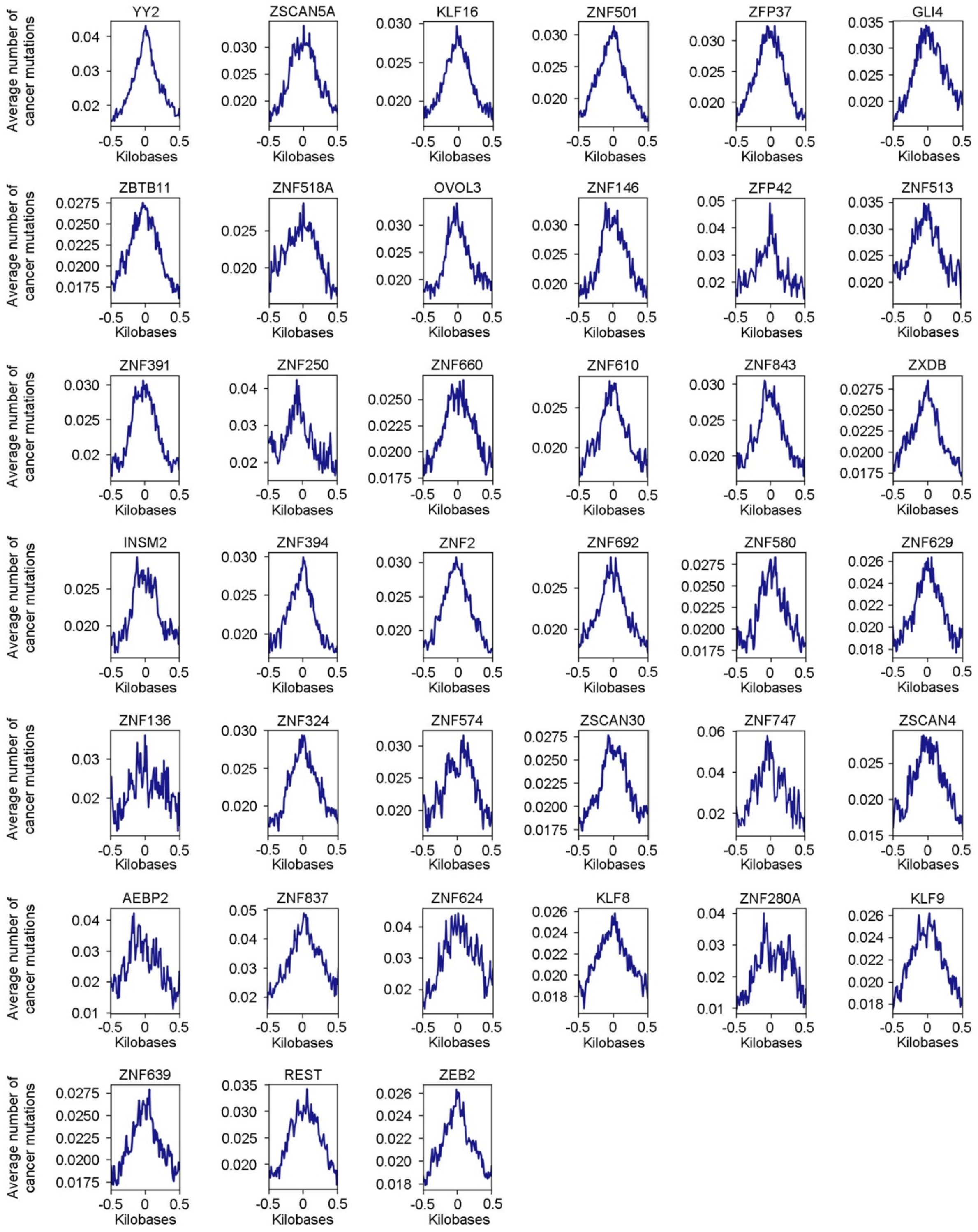

**Supplemental Figure S6. Metaplot representation of cancer mutations around DNA-binding sites for C2H2-ZFPs at promoter LRI anchors +/- 500 bp**. The figure only displays C2H2-ZFPs that overlap significantly with cancer mutations with all three different backgrounds used in Fig. 5A and that were not included in Fig. 5D.

Supplemental Figure 7

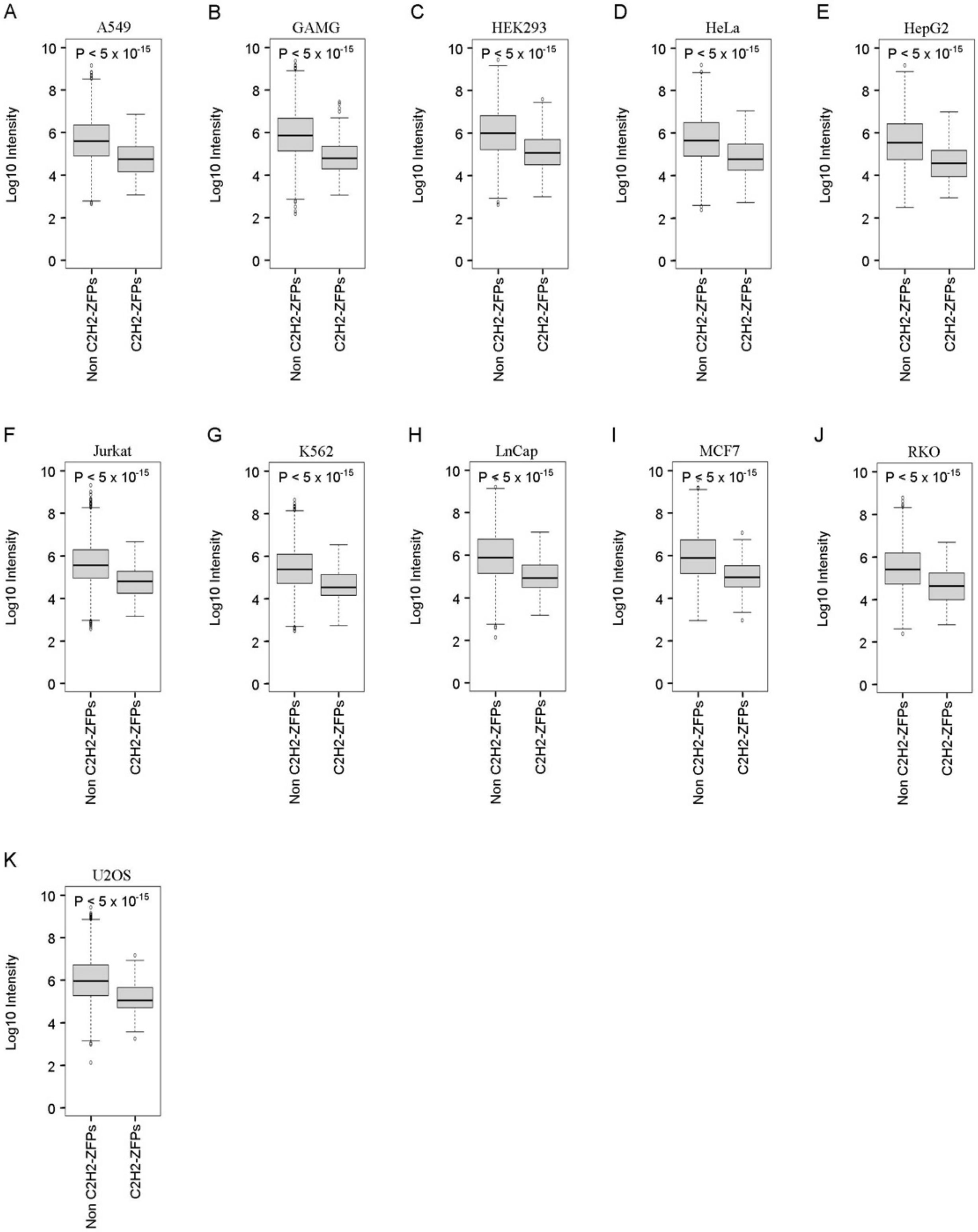

**Supplemental Figure S7. C2H2-ZFPs are expressed at lower levels compared to non-C2H2-ZFP proteins in the human proteome.**

(A-K) Boxplots comparing the expression of C2H2-ZFPs to non-C2H2-ZFP proteins in (A) A549, (B) GAMG, (C) HEK293, (D) HeLa, (E) HepG2, (F) Jurkat, (G) K562, (H) LnCap, (I) MCF7, (J) RKO, and (K) U2OS cell lines. The Log10 intensity scores after iBAQ normalization are used as a quantification of protein expression. The p-values were calculated by using the Mann-Whitney test.

### Supplemental Figure 8

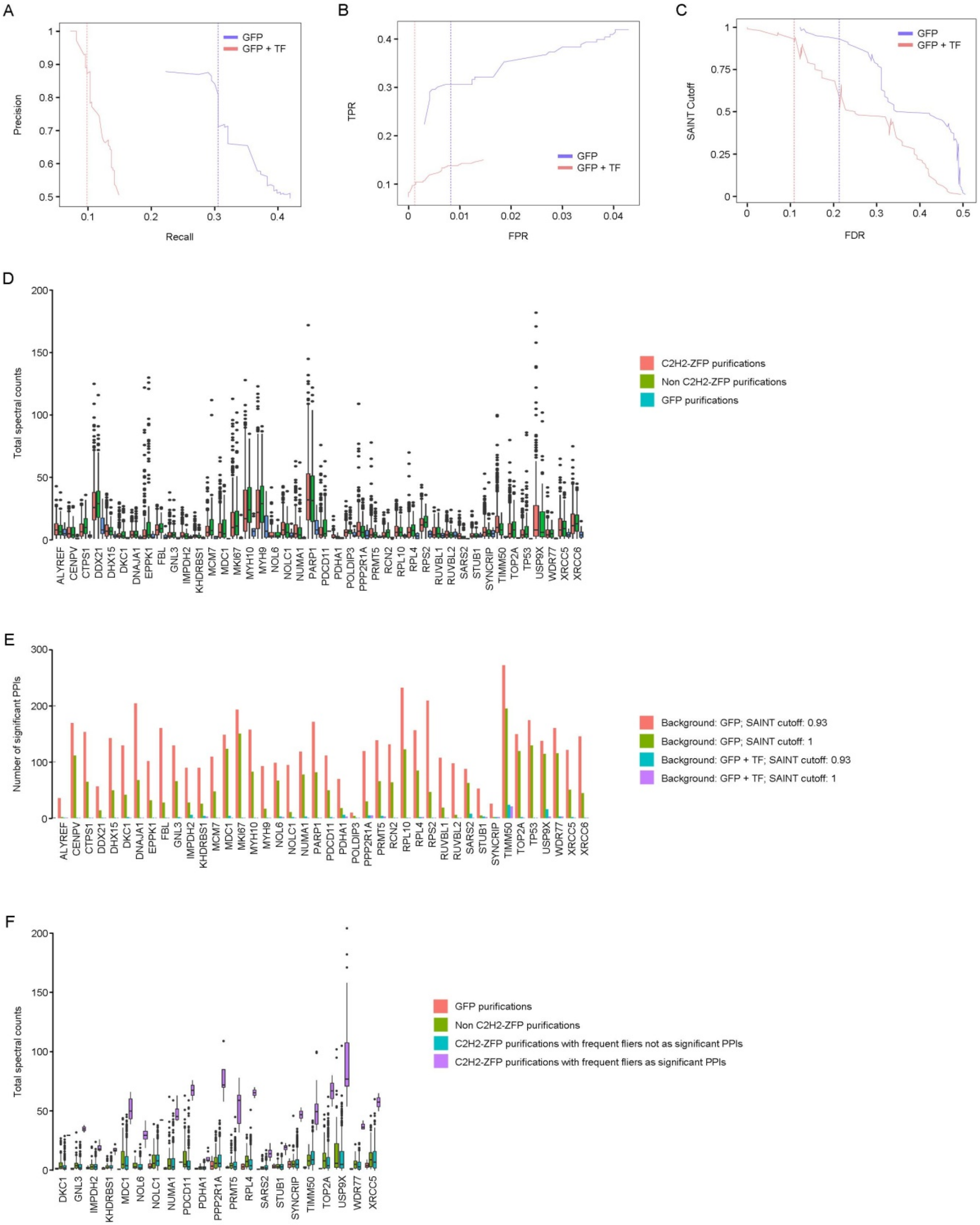

**Supplemental Figure S8. Assessment of scoring method for C2H2-ZFP PPIs.**

(A) Precision-recall curves comparing SAINT analyses using GFP only purifications (GFP) or GFP and non-C2H2-ZFP TFs as background (GFP + TF). Vertical lines indicate a SAINT cutoff of 0.93 for the GFP and GFP + TF curves.

(B) ROC curves comparing SAINT analyses using GFP only purifications or GFP + TF as background. Vertical lines indicate SAINT cutoff of 0.93 for the GFP and GFP + TF curves.

(C) Curves comparing FDR at different SAINT cutoffs using GFP only purifications, or GFP + TF as background. Vertical lines indicate SAINT cutoff of 0.93 for the GFP and GFP + TF curves.

(D) Boxplots showing the distributions of total spectral counts of some proteins that appear frequently in AP/MS experiments in GFP purifications, C2H2-ZFP purifications, and non-C2H2-ZFP TF purifications.

(E) Bargraph showing the number of significant PPIs involving some proteins that appear frequently in AP/MS experiments after SAINT scoring with GFP purifications or GFP + TF as background at SAINT cutoff of 0.93 and 1.

(F) Boxplots comparing the distributions of total spectral counts for proteins that appear frequently in AP/MS experiments that significantly interact with a subset of C2H2-ZFPs (SAINT score ≥ 0.93) to purifications done for GFP, non C2H2-ZFP TFs and C2H2-ZFPs that do not interact significantly with those proteins.
